# Integrating molecular and physiological approaches to quantify genetic controls for wheat development and improve phenotyping

**DOI:** 10.1101/2025.09.11.675709

**Authors:** Hamish Brown, John McCallum, Paul Johnston, Meeghan Pither-Joyce, Richard Macknight, Neil Huth, Derrick Moot, Bangyou Zheng, Zhigan Zhao, Enli Wang

## Abstract

Disentangling genotype × environment (G×E) effects is critical to understand the performance of wheat across different environments. A framework for doing this was previously presented in a model that integrated knowledge of crop physiology and the *Vrn* gene feedback loop to explain and predict the time of anthesis. The aims of this study were: 1) provide an updated description of the Cereal Anthesis Molecular Phenology (CAMP) model; 2) to verify the model’s assumptions regarding the relationship between *Vrn* gene expression and the timing of phenological stages in a set of diverse genotypes and environments; 3) to use the CAMP model to establish a phenotyping strategy for use in genetic studies and model parameterisation.

Six wheat genotypes with a range of cool temperature and photoperiod sensitivities were evaluated. Apical development, final leaf number (FLN) and temporal expression of *Vrn1, Vrn2* and *Vrn3* were compared with model predictions. There was a clear relationship between *FLN* responses to cool temperature and photoperiod, the timing of phenological events and the patterns of *Vrn* gene expression for all genotypes. There was general agreement between the temporal patterns of foliar gene expression observed with those assumed by CAMP, but some obvious discrepancies. These may be related to differences between gene expression in foliar (observed) and apical (assumed by the model) parts of the plant, or differences in the way observed and modelled gene expression are scaled. Overall, the model described all the observed development responses to environment and provides a basis for building quantitative predictions of field-based development from genotypic and environmental data. A protocol is presented for phenotyping wheat using FLN measured in specific combinations of temperature and photoperiod. It allows easy and unconfounded measure of key developmental phenotypes that clearly relate to the genetic make-up of the plants and underlying gene expression profiles.

## 1. Introduction

The key phenotype for matching cereal genotypes best suited to a given environment is anthesis date. A substantial amount of effort is invested in understanding the genetic controls of this phenotype. A measure of when anthesis occurs in days after sowing (DAS) does not provide a stable phenotype because different genotypes respond to temperature and photoperiod (Pp) in different ways, so DAS interacts with location and sowing date. Process models are useful tools for quantifying genotype by environment by management (GxExM) responses and providing genotype-specific coefficients that can be used as stable phenotypes giving more power in genomic analyses (Wang *et al*., 2019). The ‘Kirby framework’ (Brown *et al*., 2013) provides a useful structure for capturing the GxExM of anthesis timing by representing the underlying physiological mechanisms. Specifically, flag leaf appearance always precedes anthesis and prediction of its timing is critical. The time of flag leaf appearance depends on the number of main-stem leaves produced, referred to as final leaf number (FLN), and the thermal time duration that is required for each leaf to appear (called Phyllochron). Phyllochron, quantified as °Cd per leaf, changes with crop ontology, photoperiod and genotype (Slafer & Rawson, 1997; Baumont *et al*., 2019; Bloomfield *et al*., 2023).

The FLN of a wheat crop depends on the prevailing environmental conditions and the genotype’s sensitivity to photoperiod and cool temperatures (Brooking *et al*., 1995; Robertson *et al*., 1996; Brooking & Jamieson, 2002; Gonzalez *et al*., 2002; Bloomfield *et al*., 2023). A complete quantitative understanding of the response of FLN to genotype and environment is still lacking. Brown *et al*., (2013) attempted to progress this understanding, proposing an integrated model, combining molecular pathway knowledge of development processes with a deterministic phenology model. Herein we refer to the conceptual model described by Brown *et al*., (2013) as the Cereal Anthesis Molecular Phenology (CAMP) model. The CAMP model provides a quantitative framework in which its parameters can be considered to represent expression rates of virtual genes that may ultimately be linked to marker, single nucleotide polymorphism (SNP), QTL or genome data to enable estimation of the anthesis date of any known genotype in any defined environment (Zheng *et al*., 2025). The temporal patterns of gene expression predicted in CAMP are not of absolute concentrations, rather expression relative to an arbitrary level required to trigger an event. The algorithms for predicting their expression were developed using expected patterns of gene expression required to reach key phenological events. However, they have not been validated with experimental material where temporal patterns of gene expression and observations of phenological events have been observed simultaneously.

Here, we further improved the conceptualization of the CAMP model based on latest understanding and provide a full formal description of the model and its parameterization scheme in the supplementary material. We firstly used the mechanistic framework encapsulated in CAMP to design a set of developmental phenotypes and specify a set of environmental treatments needed to quantify these. Then we describe results from an experiment where six diverse genotypes of wheat were grown with different temperature and photoperiod treatments and their *Vrn* gene expression and phenology throughout the vegetative and early reproductive phases were measured. We finally compared these data to the predictions of CAMP to verify its concepts of gene expression and genetic control.

## 2. Materials and Methods

### 2.1 CAMP model general description

A full description of the CAMP model’s algorithms is given in the supplementary material including a calculation scheme for deriving its parameters from the developmental phenotypes calculated from FLN measurements described in this paper. Here we provide an updated overview of the model first proposed by Brown *et al*., (2013) and improved as a result of the work reported in this and an allied paper (Wang *et al*., 2025).

The timing of flag leaf appearance is central to the model and CAMP splits this period into four phases:

1. An Emerging phase (Em), from seed imbibition to emergence, when exposure to lower temperature may reduce FLN.
2. A Vegetative phase (Veg), from emergence to vernal initiation (VI), when exposure to lower temperature may reduce FLN and long photoperiod (Pp) may increase or decrease FLN.
3. An Early Reproductive phase (ER), from VI to terminal spikelet (TS), when exposure to long photoperiod may reduce FLN.
4. A stem extension phase, from TS to flag leaf expanded (FLE), when FLN has been fixed and remaining leaves must emerge.

The separation of the time from emergence to TS into two phases is a critical feature of the model. The wheat crop exhibits vernal (spring like) behaviour in ER phase where the crop accelerates development in response to lengthening photoperiod. The wheat plant also responds to photoperiod in the phase from emergence to VI, but the response can be in the opposite direction, so that it is essential to separate these phases so environmental response and genetic controls can be effectively disentangled.

With the importance of the VI and TS stages established, the CAMP model then uses predictions of Vrn gene expression to predict their timing. Three key genes control the developmental progression towards TS. *Vrn1* is a meristem identity gene that triggers the differentiation of spikelets on the apex when expressed at sufficiently high levels (Trevaskis, 2010). *Vrn3* is the florigen gene and promotes Vrn1 to levels needed to start spikelet differentiation (Yan *et al*., 2006). *Vrn2* blocks the action of Vrn3 and is downregulated by *Vrn1* (Li & Dubcovsky, 2008), so when present, creates a requirement for greater expression of *Vrn1* before Vrn3 may promote progress toward VI. *Vrn3* may be further upregulated in the ER phase to accelerate progress to TS (Yan *et al*., 2006). *Vrn1* may be upregulated by exposure to cold temperatures and *Vrn2* and *Vrn3* are upregulated by exposure to longer photoperiods.

To reach VI, the CAMP model assumes *Vrn1* must first be expressed to sufficient levels to block any Vrn2 and allow *Vrn3* expression. Then the combined promotional effects of Vrn1 and Vrn3 must exceed an arbitrary threshold to trigger VI. *Vrn1* is expressed at different base levels depending on genotype and may be upregulated in response to cool temperatures to accelerate progress toward VI. *Vrn2* is expressed at different levels depending on genotype and is upregulated by long photoperiod such that exposure to short photoperiod accelerates progress toward VI. Once *Vrn2* is downregulated, Vrn3 can become active, upregulating *Vrn1* to facilitate spikelet differentiation. *Vrn3* is upregulated in response to long photoperiod which can accelerate development through Veg and ER stages depending on genotypes’ photoperiod sensitivity.

*Vrn1* is upregulated regardless of temperature in spring genotypes, but upregulated faster at lower temperatures in winter genotypes (Trevaskis *et al*., 2007; Diaz *et al*., 2012). As such, linking genotypic data to *Vrn1* expression coefficients gives a mechanism for quantifying low-temperature responses in the timing of VI. *Vrn2* is upregulated in response to long photoperiod in some genotypes but absent in the genome others (Yan *et al*., 2004; Distelfeld & Dubcovsky, 2010; Li *et al*., 2011). Thus, linking genotypic data to *Vrn2* expression coefficients gives a mechanism for quantifying short photoperiod responses in the timing of VI. *Vrn3* is upregulated in response to long photoperiods and the extent of response is dependent on photoperiod genes (Yan *et al*., 2006; Sasani *et al*., 2009; Kitagawa *et al*., 2012; Shaw *et al*., 2012). Therefore, linking genotype data to *Vrn3* expression coefficients gives a mechanism for quantifying long photoperiod responses in the timing of VI and TS. Linking gene expression to the timing of phenological events, and further linking phenology to FLN provides a framework to understand G×E at the molecular and physiological levels. It also allows us to design methods to quantify G×E for the different developmental phenotypes (DP) of wheat.

FLN is set when TS occurs and the crop’s phyllochron is used to predict leaf appearance (quantified by Haun Stage, HS) to determine when FLE occurs. Temperature is a key driver or to timing of FLE, independent of photoperiod and cool temperature effects on FLN, because it drives the development of HS. This effect is captured in CAMP by deriving apparent gene expression and the timing of VI and TS relative to HS rather than days. The duration from FLE to anthesis is calculated from a genotype specific thermal time target and photoperiod sensitivity.

### 2.2 Quantification of Developmental Phenotypes

#### 2.2.1 Environmental treatments

The analysis scheme developed below uses measurements of *FLN* of wheat genotypes grown with cool- or warm-temperature treatment soon after seed imbibition (C and W, respectively) in factorial combination with long and short photoperiod from emergence (L and S, respectively). This requires the definition of appropriate conditions to achieve complete and nil response to low temperature and ensure long and short photoperiods are close to the upper and lower response thresholds. To ensure photoperiod responses are not confounded by low-temperature responses and vice versa, a cool treatment should be applied as early as possible. Wheat begins to accumulate a low-temperature response as soon as the seed is imbibed, and this can be saturated by holding imbibed seed at 1°C in the dark for >90 days (Brooking & Jamieson, 2002; Dixon *et al*., 2019). The process of germination and emergence takes ∼100°C days so the cold temperature response can be saturated before emergence and these conditions are what were used to apply C treatments in this study. Alternatively, the fewest days to saturate vernalisation occurs at 6°C, so sowing seed into pots kept at this temperature can also achieve complete low-temperature response at or soon after HS = 2 (Bloomfield *et al*., 2023). Temperatures of up to 15°C have been shown to induce a reduction in FLN (Brooking & Jamieson, 2002; Ritchie *et al*., 2015), so plants in the W treatment were kept with day and night temperatures >20°C during the Em and Veg phases to ensure no cool-temperature responses occurred. For wheat, the response to photoperiod is well represented assuming a linear increase from a base of 8 h to an upper threshold of 16 h (Jamieson *et al*., 1998) so we use 8 h to represent the S treatment and 16 h to represent the L treatment.

#### 2.2.2 Minimum Leaf Number (*MinLN*)

If a genotype receives C treatment, VI will be reached at its earliest possible time and, if grown in L conditions, it will also progress to TS at the earliest possible time. Thus, in CL conditions, the observed *FLN* will represent its minimum (*MinLN*) and can be used as a measure of earliness per se for the genotype (Figure 1):

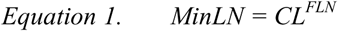

**Figure 1.**
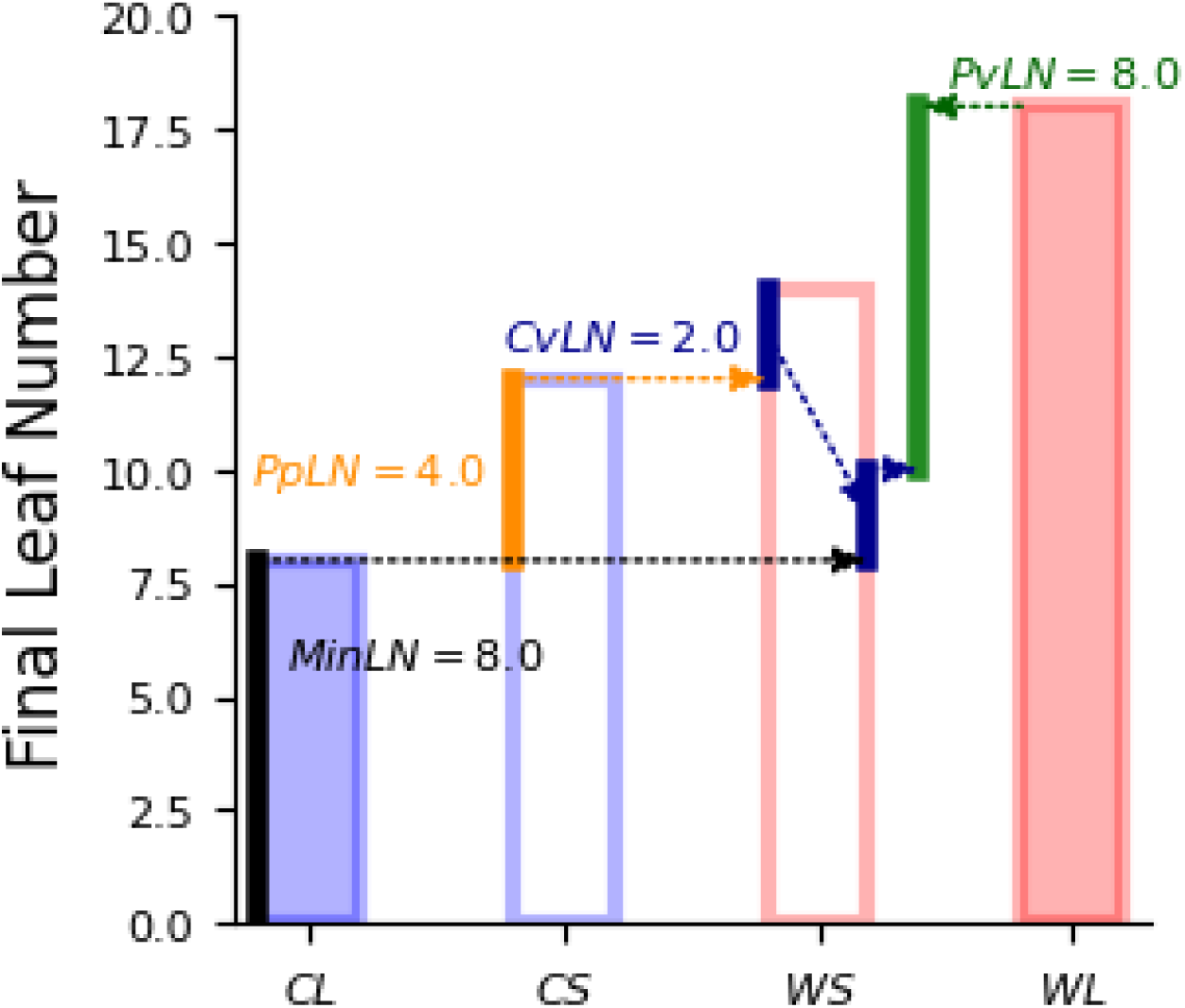
Final leaf numbers for ‘Batten’ winter wheat taken from selected treatments published by (Brooking & Jamieson, 2002) with and without cold-temperature treatment during the emerging phase (C and W, blue and red bars, respectively), then grown under long (L, filled bars) and short (S, open bars) photoperiod. The difference scheme demonstrates how minimum final leaf number (MinLN), photoperiod sensitivity (PpLN), cold vernalisation sensitivity (CvLN) and photoperiod vernalisation sensitivity (PvLN) are calculated. PvLN can be negative or positive.

#### 2.2.3 Photoperiod (Pp) sensitivity (*PpLN*)

If a genotype receives C treatment and is grown in S conditions its *FLN* may increase (Figure 1). Downregulation of *Vrn3* will extend the duration of the ER phase. Expression of *Vrn3* decreases with shortening photoperiod and the extent of this decrease depends on genotype (Yan *et al*., 2006; Sasani *et al*., 2009; Kitagawa *et al*., 2012; Shaw *et al*., 2012). When *Vrn3* expression is minimised under short photoperiod, the ER duration will reach a maximum.

Therefore, the photoperiod sensitivity of a genotype (*PpLN*) may be quantified as the difference the FLN between the L and S treatments for cold treated genotypes (Figure 1):

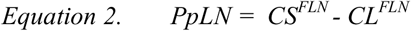

#### 2.2.4 Cold vernalisation sensitivity (*CvLN*)

When a crop is grown in continuously warm conditions (W), its Veg duration and FLN may increase. However, the extent of the *FLN* response interacts with photoperiod. If we first consider the WS treatment, *FLN* will be increased by response to short photoperiod in the ER phase but may be further extended by lack of cold exposure prior to VI. Some alleles of Vrn1 will only upregulate *Vrn1* expression in the cold (Trevaskis *et al*., 2007; Diaz *et al*., 2012). Such genotypes will reach VI more slowly if W conditions are imposed and any increase in *FLN* beyond that already attributed to *PpLN* provides a quantification of the cold vernalisation sensitivity (*CvLN*) of the genotype (Figure 1):

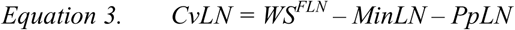

#### 2.2.5 Photoperiod vernalisation sensitivity (*PvLN*)

When a genotype is grown in warm conditions with long photoperiod (WL), the duration of its Veg phase and subsequent *FLN* may be increased or decreased relative to the WS treatment depending on the relative activity of the photoperiod sensitive genes (*Vrn2* and *Vrn3*). The photoperiod vernalization sensitivity of the vegetative phase is quantified as (Figure 1):

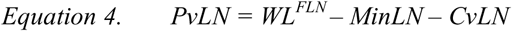

This may be a positive or negative value. In the case of a genotype that actively expresses *Vrn2* under L conditions, this will block *Vrn3* and delay VI causing *FLN* to increase and give a positive *PvLN*. This response has been described as short-day vernalisation (Brooking & Jamieson, 2002; Allard *et al*., 2012). In the case of a genotype with low *Vrn2* activity, *Vrn3* may be upregulated during the vegetative phase which will make VI earlier than under S, reducing *FLN* and giving a negative number for *PvLN*.

### 2.3 Plant material and culture

The six genotypes selected for this study span a diverse range of vernalisation and photoperiod sensitivities observed in the field. The ‘Batten’ near isogenic lines (winter and spring) have different vernalisation sensitivities (Brooking & Jamieson, 2002) while Otane is a true spring wheat with no vernalisation response. ‘Saracen’ is a cultivar with intermediate vernalisation requirement; ‘Amarok’ is a winter wheat cultivar with low photoperiod sensitivity and CRWT153 is a breeding line that consistently displays very late anthesis in field trials. The developmental phenotypes of these genotypes observed in the field were consistent with the allelic composition for *Vrn* and *Ppd* genes revealed by published markers (**Error! Reference source not found.**).

Plants were grown in a Conviron growth room (Controlled Environments Ltd., Manitoba, Canada) at the Lincoln University Biotron facility (Shi *et al*., 2011). Growth rooms were run with a constant 16-h photoperiod and 25/20°C light/dark temperature (L) or an 8-h photoperiod and 25/22°C light/dark temperature (S). Lights were switched on in three separate banks at 30 minute intervals at the beginning of the photoperiod and produced photosynthetic photo flux density of 1150 μ mol m^2^ s^-1^ when all three banks were on. Lights were switched off one bank at a time in 30 min intervals at the end of the photoperiod. Pots of three to six plants of the same genotype were established in bark and soil based potting mix. Pots were assembled on 1.2 m high benches with raised sides and a central drainage hole. Up to 600 pots could be included with this layout. The growth room had mirrored walls and a well-mixed atmosphere to provide even environmental conditions. Pots were laid out in a random block design to account for any possible variation in air flow or radiation.

For warm-temperature treatments (W), six seeds were sown in each pot and covered with a 15-mm layer of vermiculite. Vermiculite was used instead of potting mix because it offers little resistance to coleoptile extension, ensuring even emergence. It also has a high albedo which limits radiant heating of the pots to assist with keeping the apex temperature of the wheat consistent with air temperature.

For the cool-temperature treatments (C), seeds were sterilised by immersion in 4% sodium hypochlorite and stirred for 30 min. Seeds were then rinsed with reverse osmosis water and transferred to sterile Petri dishes containing moist filter paper. Petri dishes were put inside a polystyrene box to exclude light and maintain a stable temperature. This box was kept in a refrigerator (1°C +/- 0.5 °C) for 90 days and dishes were checked weekly. Re-wetting of filter paper with reverse osmosis water was undertaken as needed and any seeds showing signs of fungal growth were removed. A Hobo temperature logger (Onset Computer Corp. MA, USA) was kept in the box to record hourly temperature. At the completion of this treatment, seeds had germinated with a 1- to 5-mm coleoptile and 10- to 20-mm roots visible. These plants were carefully teased apart, transplanted into 1-L pots (3–6 plants per pot) and covered with a 15mm layer of vermiculite (as for the W treatments). From this time on plants were grown in the same conditions as the W treatment.

Pots were watered with a watering can until one week after emergence. At this time an irrigation dripper was installed in each pot and moisture sensors installed in four sentinel pots (pots with well-established plants). An irrigation control system using a CR1000 data logger (Campbell Scientific, Logan, UT, USA) turned on all drippers for 1 min each time the mean water content of the sentinel pots reached a specified trigger point. Any water draining from the tables was plumbed into the growth chamber’s condensation collection system. This method enabled the continual irrigation of many pots without wetting the foliage of the plants so avoiding foliar disease problems. Beyond eight weeks, plants showed evidence of nutrient stress which suggests their growth demands had exceeded the supply of nutrients in the potting mix. From this point, each pot received 100 mL of Hoaglands (Hoagland and Arnon 1950) solution weekly.

### 2.4 Plant sampling

Samples were taken at weekly intervals. The timing of the light period was set so the lights went out at 10:00 am. This allowed time to take leaf samples for RNA analysis at the end of the light period when gene expression was determined to be most representative of the treatments (see Supplement B). For RNA isolation, the most recent fully expanded leaf was cut (using sharp scissors) at its mid length and immediately, ∼5 × 1-mm strips (∼1 cm^2^, ∼25 mg) were cut from the basal end of the sampled leaf directly into a 1-mL tube containing 300 µL of DNA/RNA Shield (Zymo Research Ltd.). Care was taken to completely submerge the leaf tissue in the DNA/RNA shield. Samples were stored at −20^°^C for later RNA isolation. One pot was taken from each replicate (three pots per treatment) giving 9–18 plants per treatment for dissection at each sampling date. Plants were removed from their pots, the number of fully expanded main-stem leaves (those with ligules visible) were counted, and the length of the most recently fully expanded leaf and the current expanding leaf was measured for retrospective calculation of HS (Brooking & Jamieson, 2002). The irrigation method meant the canopies remained dry and senesced leaves did not decompose, allowing the number of emerged leaves to be determined accurately. Each plant was then dissected under a binocular microscope to observe and count the number of un-emerged leaves and the number of primordia on the apex that were yet to differentiate and emerge (Brooking & Jamieson, 2002). Measurements were continued until seed heads emerged. At this time, any remaining pots (6–12 pots giving 30–60 plants per treatment) were destructively sampled to measure *FLN* and the number of spikelets on the spike.

### 2.5 Observed timings of Vernal Induction and Terminal Spikelet stages

The timings of VI and TS were quantified in terms of the Haun Stage (HS) at which they occurred and are superscripted with HS to show this. During the Veg phase, wheat produces primordia at a rate about double that of ligule appearance (Brooking & Jamieson, 2002). This rate increases when the crop reaches *VI* and remains elevated until *TS*. As such the HS timings of these events can be identified as break points on a plot of total organ number (leaves, undifferentiated primordia and spikelets) against HS at each destructive sampling.

### 2.6 RNA isolation and quantification

Frozen samples were allowed to equilibrate to room temperature. A leaf homogenate was made by adding 0.5-mm diameter glass beads to the sample tube and grinding with the flat end of a 0.5-mL microtube pestle. After centrifuging at 2000 × g for 10 min, 60 µL of supernatant was transferred to a new 0.5-mL deep well plate and an equal volume of RNA Lysis Buffer (Zymo Research) was added. Total RNA was isolated using the ZR-96 Quick RNA Kit (Zymo Research) according to manufacturer’s instructions and serially eluted with 1 × 25 µL and 1 × 10 µL DNase/RNase free water preheated to 95°C. RNA concentration was determined using a Victor^3^ Multilabel Plate Reader (Perkin Elmer) and the Quant-IT RiboGreen RNA quantification kit (Invitrogen).

### 2.7 Two-step quantitative real-time PCR (RT-qPCR)

Reverse transcription was carried out with 100 ng of total RNA in a volume of 20 μL using Maxima First Strand cDNA Synthesis Kit for RT-qPCR, with dsDNase (Thermo Scientific) according to manufacturer’s instructions. A replicate template plate for each primer set was made by diluting first-strand cDNA 5-fold with double distilled H_2_O and transferring 3 × 5- µL aliquots (technical replicates) into PCR plates. An identical sample was loaded into every template plate as an Inter-Run Calibrator (IRC) and plates were stored at −80°C. RT-qPCR was performed in 10-µL reaction volumes in 96-well plates on the StepOnePlus Real-Time PCR System (Applied Biosystems) for Experiment 1, or in 384-well plates on the LightCycler 480 System (Roche Life Science) for Experiment 2, using SYBR Green detection and Luminaris Color HiGreen High ROX qPCR Master Mix (Thermo Scientific) according to manufacturer’s instructions. The cycling conditions were: 2 min hold at 50°C, 10 min hold at 95°C, followed by 40 amplification cycles of 15 sec at 95°C, 1 min hold at 60°C. After completion of qPCR, melt curve analysis was carried out on the amplicons using the default settings of the Real-Time PCR instrument.

### 2.8 Primer set validation

Initially, published primer sets for *Vrn1* (WAP1), *Vrn2* (ZCCT1), *Vrn3* (WFT) and house- keeping genes (HKG) *ubiquitin* (Shimada *et al*., 2009) and *Ta54227* (Paolacci *et al*., 2009) were validated. This was done using qPCR with six serial, 4-fold dilutions of pooled cDNA (1:5), followed by melt curve analysis, as per MIQE guidelines (Bustin *et al*., 2009). qPCR efficiency was calculated for each primer pair from a standard curve using StepOne Software (Applied Biosystems). The *Vrn2* primer failed because it only had a 0.5°C difference between the primer melting temperature (Tm) and hairpin temperatures and it amplified genomic DNA. The *Vrn3* primer failed because it amplified genomic DNA and the Ubiquitin primer failed because of very low efficiency. As alternatives, primers for *Vrn2* and *Vrn3* were designed using Primer3 (Untergasser *et al*., 2012) to span exon-exon junctions, produce amplicons between 130 to 180 bp, Tm around 60°C, and low dimer and hairpin Tm. Other published primers for *Vrn2* (Dubcovsky *et al*., 2006), *Vrn3* (Yan *et al*., 2006) and the HKG, *Tef-1*α, (Giménez *et al*., 2011) were also validated and the best-performing sets selected. A description of each of the primers used in this study is given in the supplementary material (Table SA1).

### 2.9 Verification of CAMP predictions

#### 2.9.1 Model set-up and operation

The CAMP model was coded into a Python script which is available at https://github.com/HamishBrownPFR/CAMP/blob/master/CAMP.ipynb. A formal description of the code and parameterisation scheme is given in the supplementary material. The FLN developmental phenotypes measured for each genotype (Section 3.1) were used to derive the *Vrn* expression parameters needed for CAMP. Each of the treatments was simulated using CAMP with its corresponding daily temperature and Pp, so its predictions of *Vrn* gene expression could be compared with those observed. The script running the CAMP code and producing the graphs displayed in this paper can be viewed at https://github.com/HamishBrownPFR/CAMP/blob/master/Tests/CAMPCETests.py.

#### 2.9.2 Comparing CAMP predictions with observed ***Vrn*** gene expression

Each observed qPCR value for expression of each *Vrn* gene (qpcr_vi_) was normalized relative to ubiquitin expression (qpcr_ui_) for the same sample to provide relative levels of expression:

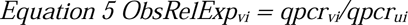

To facilitate comparisons of the relative activity of individual genes (v), they were further normalized relative to maximum expression for v to give their relative activity (*ObsRelAct_vi_*) on a scale from 0-1:

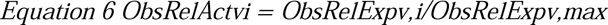

Where *ObsRelExp_v_*_,i_ represents each observation (i) of *ObsRelExp_v_* and *ObsRelExp_v_*_,max_ is the maximum observed value of *ObsRelExp_v_*.

The Camp Model predicts *Vrn* gene expression in relative amounts that are not directly comparable with *ObsRelAct_v_*. To facilitate comparison daily predictions of *Vrn* gene activity, (*camp_v_*) were normalized relative to the maximum expression predicted across all treatments (*camp_v_*_,*max*_):

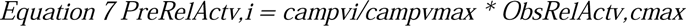

Where *ObsRelAct_v_*_,cmax_ is the maximum observed relative activity for the genotype (c) under consideration. This multiplication is included to scale the predicted expression over the same range as the observations to further facilitate comparisons. Note for *Vrn1 camp_v_* is the sum of persistent cold *Vrn1* and *VrnB* (base *Vrn1* expression). We stress that this does not provide validation of the CAMP model as the data are not sufficiently independent from the model, but also that CAMP is not coded to predict absolute gene expression. Rather, it predicts relative accumulated effects of gene activity on developmental progress. While this is not directly comparable with observations of gene expression, comparison of modelled (Equation 7) and observed (Equation 6) time courses of relative activity are valid for assessing the mechanisms assumed in CAMP relative to temporal gene expression.

## 3. Results

### 3.1 Genotypic differences in developmental phenotype

The *FLN* values recorded under each environment treatment are shown in Figure 2 and the developmental phenotypes derived for each genotype are displayed in Table 2. There was a broad range in *MinLN* from 6.65 leaves for Otane up to 14.27 leaves for CRWT153. *PpLn* ranged from no significant response in CRWT153 to a value of ∼3.8 leaves for the Batten pair. Of the winter types ‘Amarok’ had a significant *CvLN* of 5.02 but no significant *PvLN*. Alternatively, ‘Batten’ winter had no significant *CvLN* but a *PvLN* of 5.75.

**Figure 2.**
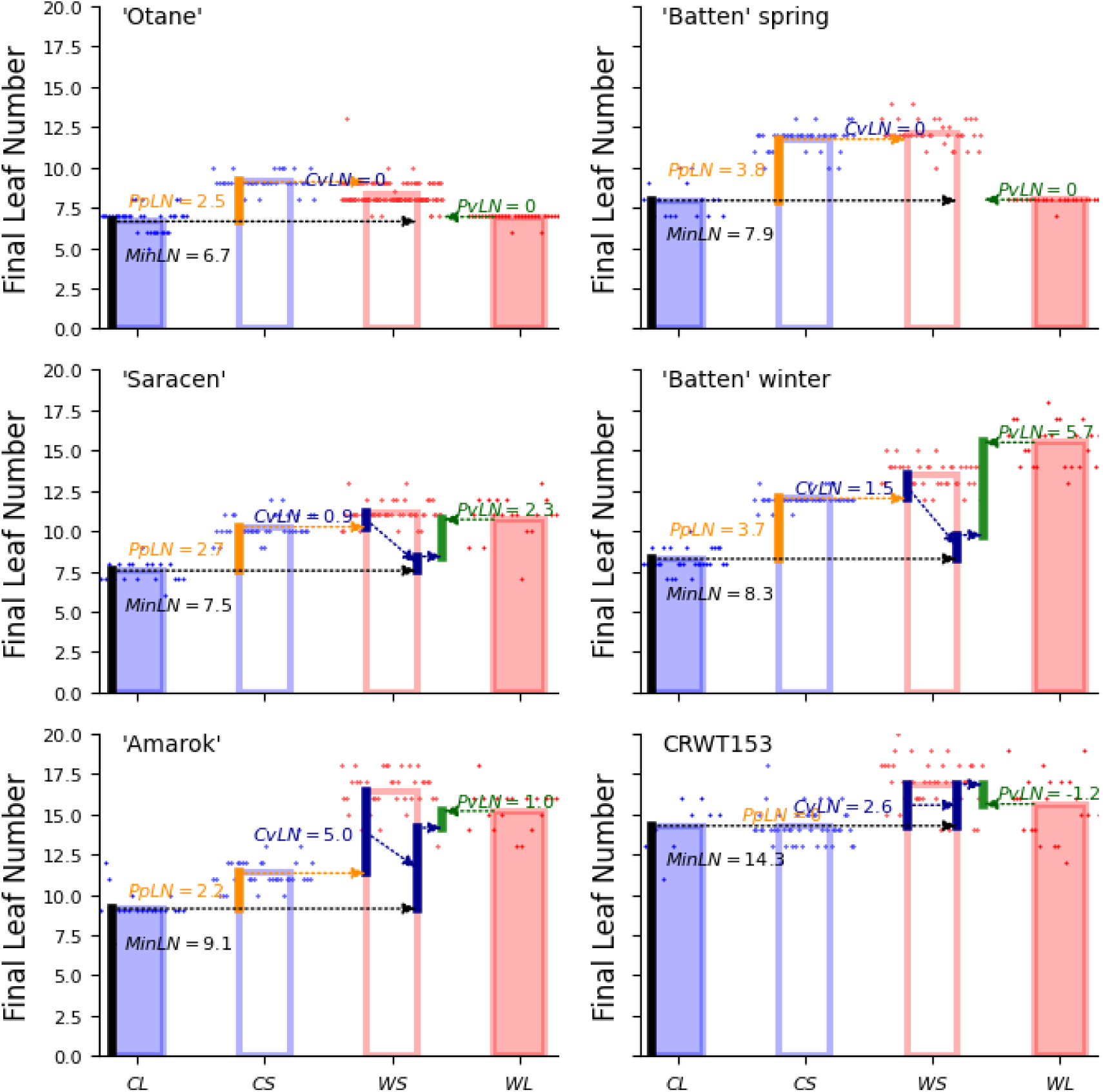
Final leaf number of different wheat cultivars grown in controlled environment under long (solid fills) or short (hollow fills) photoperiod in combination with (blue) or without (red) cool temperature treatment during the emerging phase. Points show individual observations and bars show treatment means. MinLN = minimum final leaf number; CvLN = cold vernalisation sensitivity; PpLN = photoperiod sensitivity; PvLN= photoperiod vernalisation sensitivity.

### 3.2 The Haun stage timing of key phenological events

Figure 3 shows the total organ number (TON) against HS for each treatment. The HS where observed TON deviates from the basic vegetative rate represents observed VI^HS^ and the HS where observed TON reaches a maximum represents TS^HS^. The positions of the observed and derived VI^HS^ and TS^HS^ correspond closely for all treatments providing verification for the scheme proposed for deriving these values (Supplementary material).

**Figure 3.**
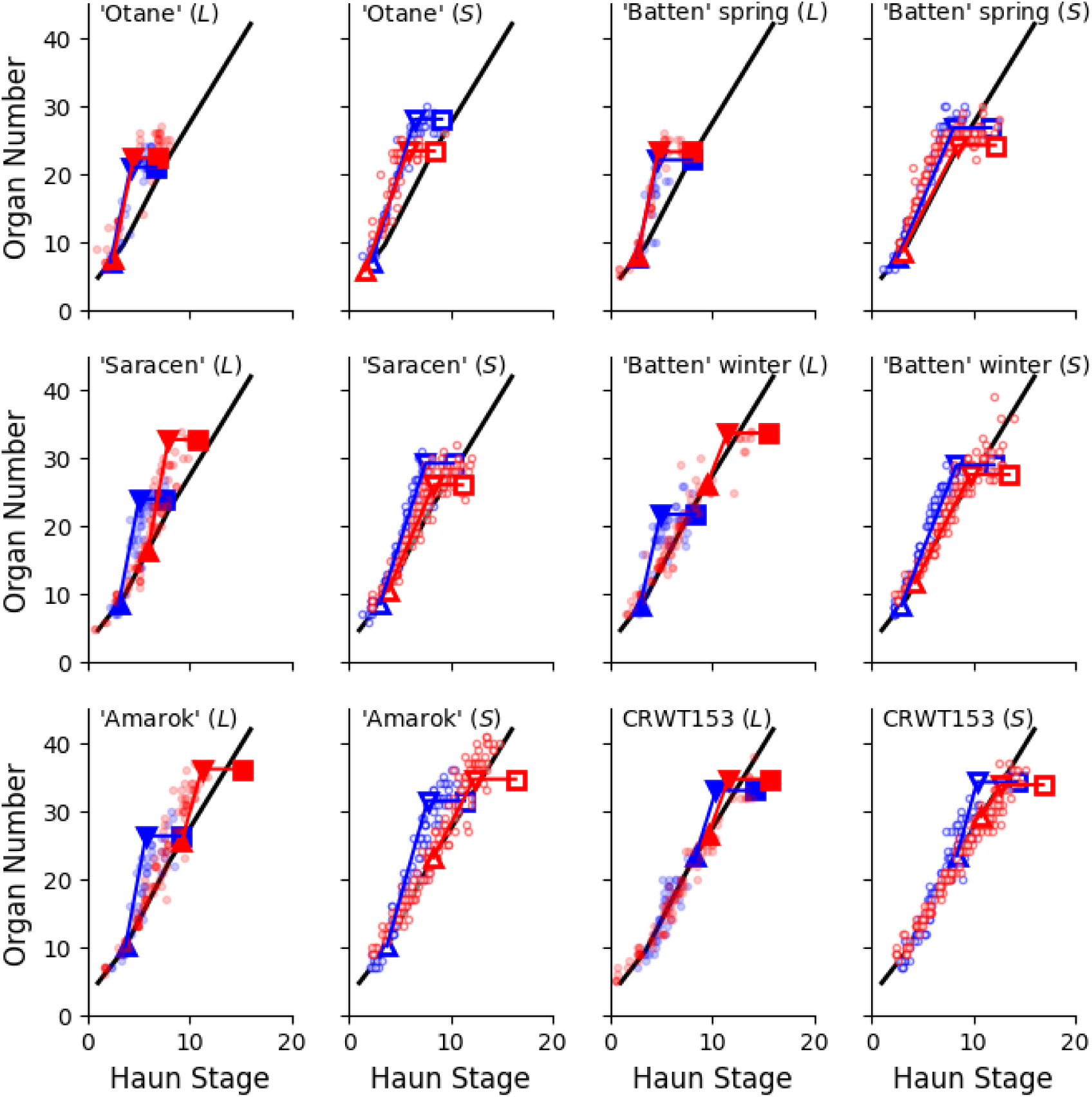
The relationship between total main-stem organ number (leaves, un-differentiated primordia and spikelets) and Haun stage (measure of main-stem leaf appearance) for different genotypes of wheat grown with (blue) or without (red) cold treatment in combination with long (filled markers) or short (open markers) photoperiod. The large squares are the organ number observed at final sampling (y-axis), plotted against observed FLN (x-axis). The large downward pointing triangles have the same y-value plotted against the TS^HS^ predicted by the CAMP parameter scheme using the observed FLN. The large upward pointing triangles are the VI^HS^ calculated from the CAMP parameter scheme (x-value) plotted against the TON produced at the base vegetative rate (displayed by the black line) at VI^HS^. The small circles represent observations of total organ number at different HS from repeat destructive samplings. The black line represents the relationship that vegetative plants are expected to conform to (Brown *et al*., 2013)

From this analysis we can compare the duration to *VI^HS^*and the duration from *VI^HS^* to *TS^HS^* for different treatments (Figure 4). For the C treatments, *VI^HS^* = 2.0 for the two spring genotypes, 3.0 for ‘Saracen’ and Batten winter, 4.0 for Amarok and 8.0 for CRWT153 and this was unaffected by photoperiod treatment. Under W treatments the *VI^HS^* of the spring genotypes was the same as under the C treatments, confirming the assumption that spring cultivars express *Vrn1* at high rates regardless of cold treatment. However, the *VI^HS^*of the winter genotypes were delayed with W treatments and there was an interaction with photoperiod as to the extent of this (Figure 4). Specifically, Batten winter showed the largest delay with *VI^HS^* of 10 for WL but a much smaller delay with a *VI^HS^* of 4.0 for the WS treatment, indicating shorter photoperiod causing less delay. Amarok also showed a similar pattern, with substantial delay in *VI^HS^* to 9.0 under WL and 8.0 under WS conditions. CRWT153 had a slight delay for the W treatments, but with slightly more delay under long photoperiod than under shorter photoperiod, a pattern different from other genotypes. ‘Saracen’ also showed a delay in *VI^HS^* to 6.0 under WL and 4.0 under the WS.

**Figure 4.**
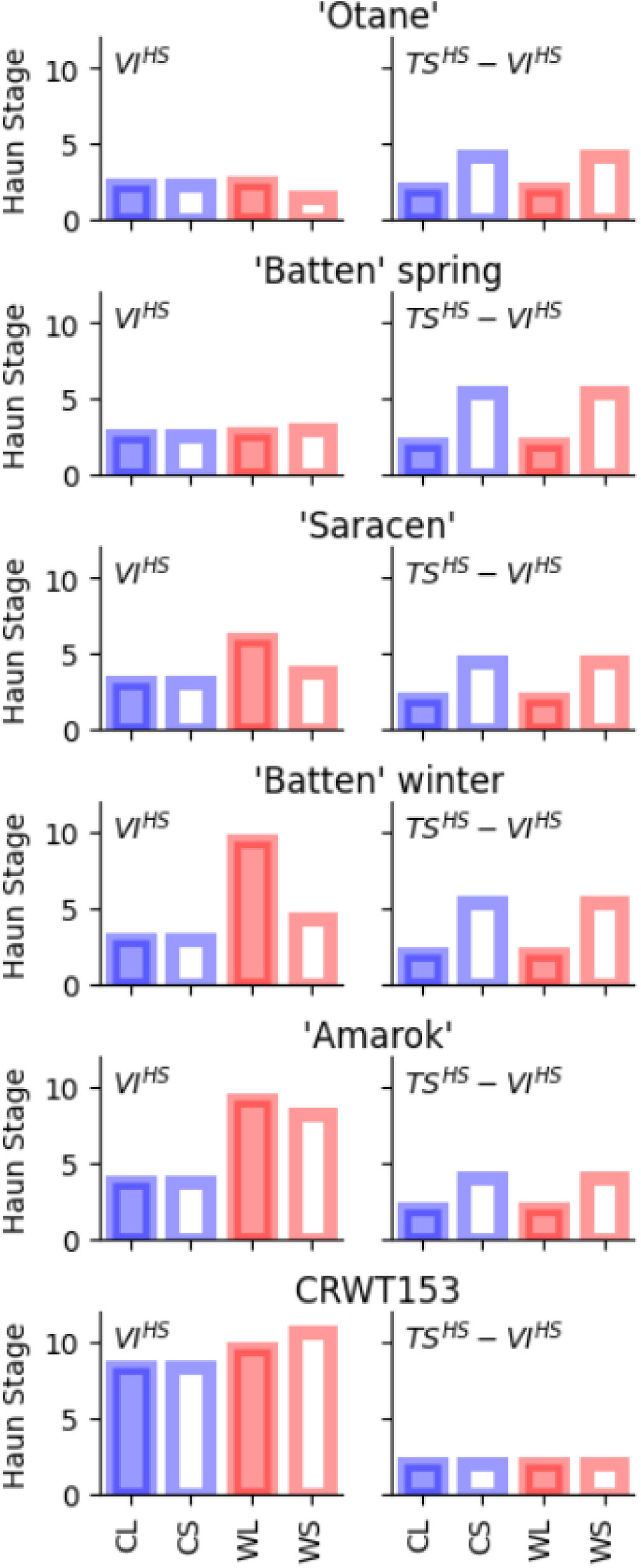
The Haun Stage timing of Vernalisation Induction (VI^HS^) and the duration of the Early Reproductive phase (VI^HS^–TS^HS^) of different cultivars grow in controlled environment under long (solid fills) or short (hollow fills) photoperiod with (blue) and without (red) cool temperature treatment of 100 days at 1°C prior to imposing photoperiod treatments.

For the duration from *VI^HS^* to *TS^HS^*, all the genotypes responded to photoperiod similarly, except CRWT153 which had no response to photoperiod (Figure 4). Regardless of C or W treatments, the duration was reduced to minimum under L and maximised under S. ‘Batten’ winter and spring had the same photoperiod response, consistent to their allelic variations (Table S1). Otane and Amarok also had very similar response to photoperiod. These results verify the assumption that response to cold temperature ended at VI.

### 3.3 *Observed Vrn* gene expression over the ontology of different genotypes in different environments

If we first consider observed expression for the CL treatments (Figure 5), we see the earliest genotypes (Otane, Batten spring and ‘Saracen’) started with *Vrn1* upregulated in their leaves, which is soon followed by upregulation of *Vrn3* and further upregulation of *Vrn1*. As we consider progressively later genotypes, the expression of *Vrn2* extends for increasing periods before upregulation of *Vrn1*, downregulation of *Vrn2*, upregulation of *Vrn3* and further promotion of *Vrn1*. In the case of CRWT153, there is limited upregulation of *Vrn1* and *Vrn3* and protracted upregulation of *Vrn2*.

**Figure 5.**
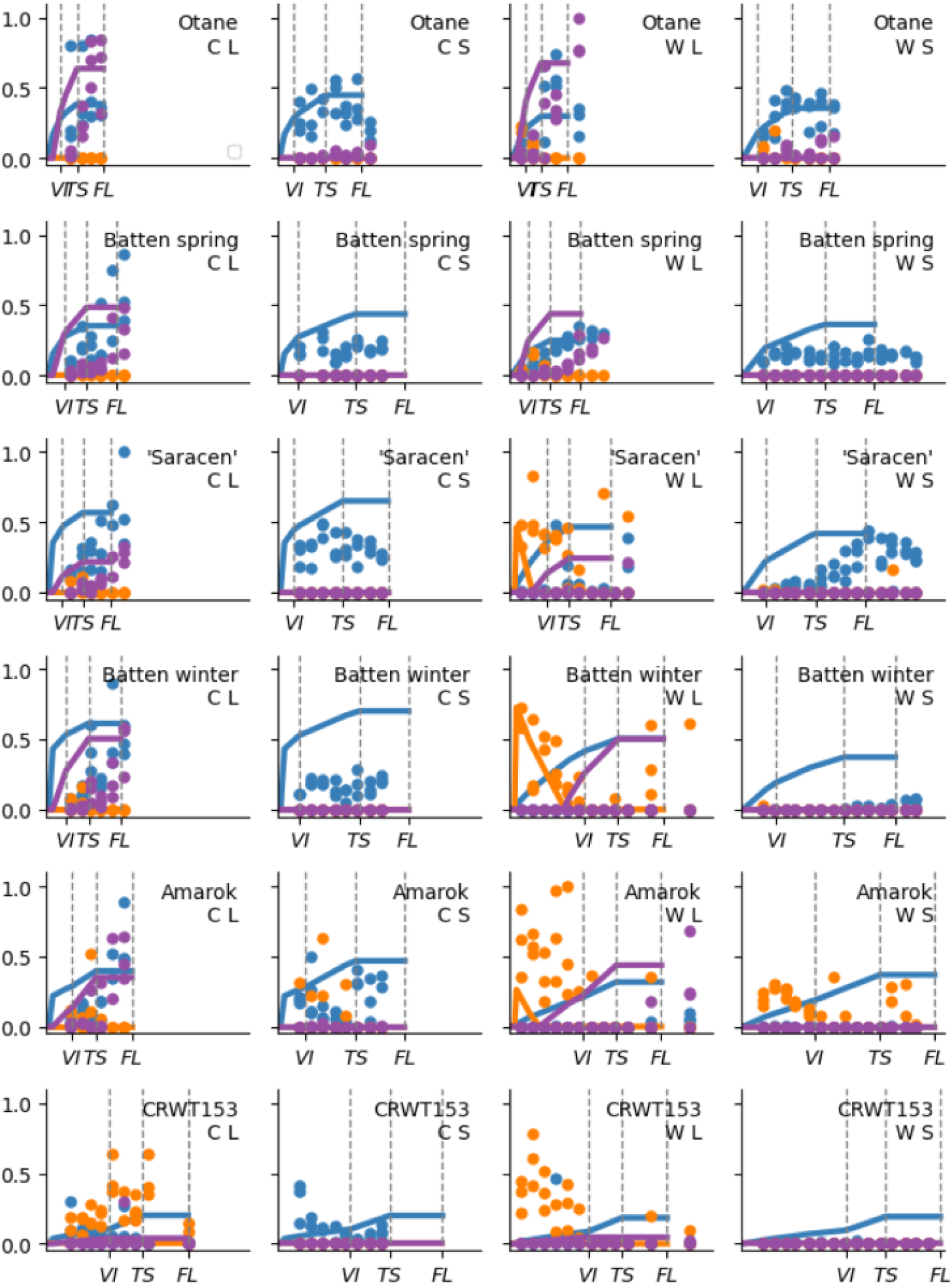
Relative expression of *Vrn* genes in six different genotypes (sorted from lowest to highest MinLN) of wheat grown under different photoperiod (S = 8 and L = 16 h, respectively) with and without cold treatment applied during the emerging phase (C and W respectively). VI, TS and FL tags show the time of vernalisation induction, terminal spikelet and flag leaf ligule appearance, respectively. The x-axis is accumulated thermal time (T_b_ = 0°C) up to a maximum of 2550 °Cd. Lines represent activity predicted by the CAMP model and circles represent corresponding observations for *Vrn1* (blue), *Vrn2* (orange) and *Vrn3* (purple).

If we next consider the WL treatments (Figure 5), we see the spring genotypes (Otane and Batten spring) quickly upregulate *Vrn1*, downregulate *Vrn2*, upregulate *Vrn3* and further upregulate *Vrn1* much the same as in the CL treatment. However, the winter genotypes show prolonged upregulation of *Vrn2* when no cold treatment is applied. The time of VI^HS^ corresponds to the downregulation of *Vrn2* but most of the winter genotypes do not display substantial upregulation of *Vrn1* or *Vrn3*.

Next, we consider the CS treatments (Figure 5) where *Vrn1* was upregulated throughout development but remained at relatively static levels. Finally we consider the WS treatments Figure 5), where we see *Vrn1* is upregulated throughout for the spring genotypes, but only upregulated after *VI^HS^* for ‘Saracen’ or not upregulated at all for the winter genotypes.

### 3.4 Comparison of model predictions

Figure 5 provides a comparison of the time course of gene activity predicted by CAMP relative to that observed for each of the six genotypes grown in each of the environmental treatments. The observed patterns of gene expressions were in general agreement with what is conceptualized in the CAMP model where: Cool temperatures upregulate *Vrn1* in winter genotypes, but *Vrn1* is upregulated in spring genotypes regardless of temperature treatment; *Vrn2* expression was limited to situations where *Vrn1* expression was low under long photoperiods; *Vrn3* expression was limited to situations where *Vrn2* expression was low under long photoperiods. However, there are also situations where the observed and predicted *Vrn* expressions differ. For example, after *VI* when *Vrn2* is supressed, CAMP assumes that genotypes should display upregulation of *Vrn3* under long Pp, but this is not reflected in many of the observed patterns (Figure 5).

CAMP tends to predict an up-regulation of relative *Vrn1* expression earlier than was observed (Figure 5). Under S conditions it predicts a continued upregulation of *Vrn1* where observed values of *Vrn1* remained static. For winter genotypes in W treatments, the model predicts an upregulation of *Vrn1* (VrnB, see supplement) where none or little was recorded. A similar pattern was evident for *Vrn3* where CAMP predicted its upregulation earlier than observed and predicted substantial upregulation for winter genotypes under W treatments when none was observed. By contrast, CAMP tended to predict *Vrn2* downregulation too early and observations of *Vrn2* were often made when the model predicted activity would be nil.

There was very good agreement between the FLN observed in each treatment and those predicted by the CAMP model (Figure 6). This should not be considered a validation of the model as the observed data have been used in deriving and testing the model. However, it provides important verification that the phenotyping scheme, method for deriving CAMP *Vrn* expression rates and the numeric implementation of the conceptual model can reproduce the broad range of FLN values created with a diverse set of genotypes and environmental combinations. In short, the phenotyping scheme and CAMP model provide a valid framework for quantifying genotypic differences and estimating how these will interact with environment to influence anthesis behaviour.

**Figure 6.**
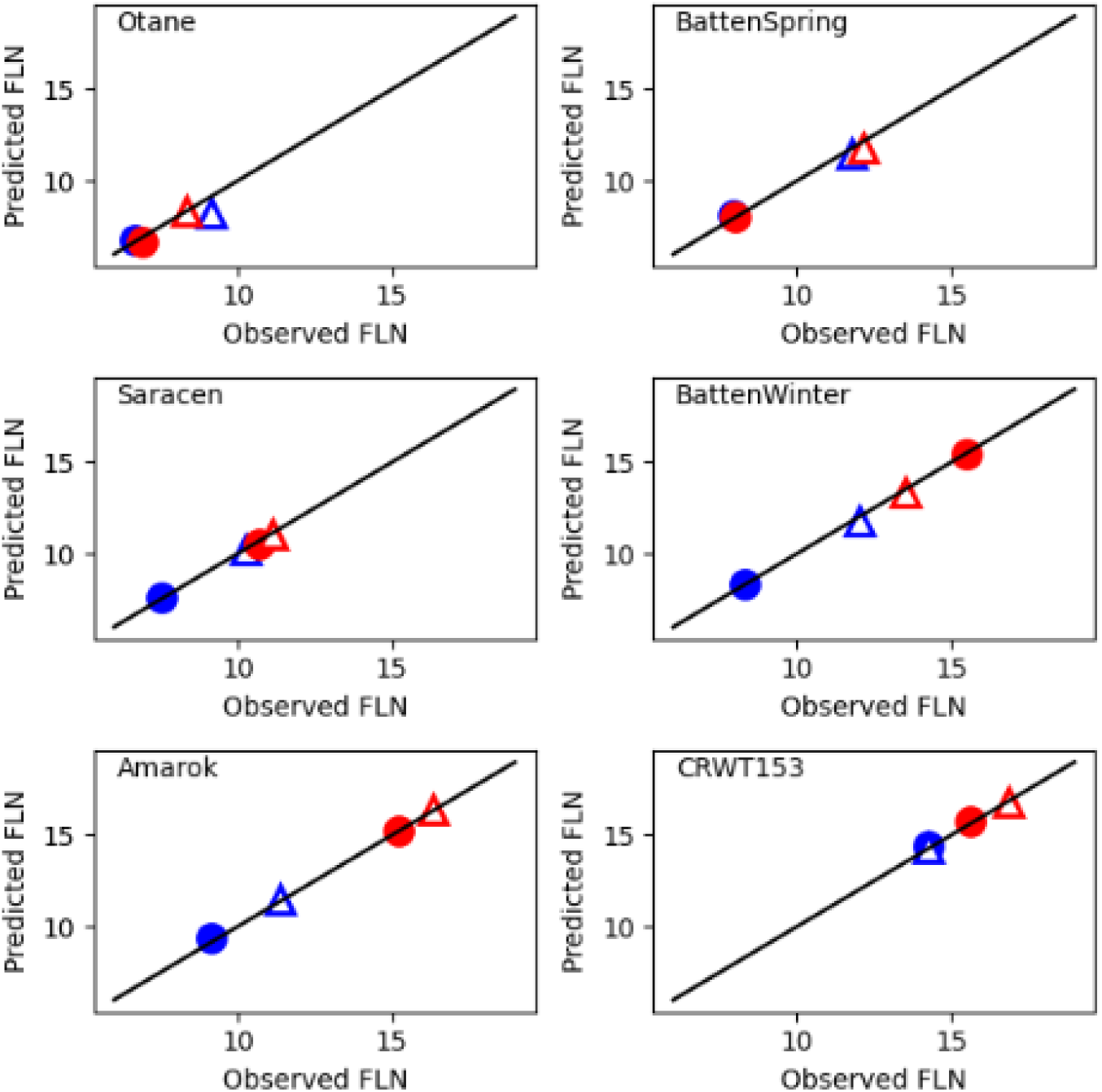
Comparison of Observed and Predicted FLN for six diverse genotypes of wheat grown under 8 (open triangles) and 16 h pp (closed circles) with (blue) and without (red) cool temperature treatment during the emerging phase.

## 4. Discussion

### 4.1 Phenotyping method

The FLN-based phenotyping method proposed in this paper (Section 2.2) provides a useful approach for characterising the developmental responses of wheat genotypes and providing data for determining parameters for the CAMP model for predictions of cereal phenology. It is expected that these phenotypes will be stable across different environments and so provide useful traits for genetic analysis of anthesis. The *MinLN* phenotype provides a quantitative representation of the genotypes earliness *per se,* which has a substantial range measured in the genotypes tested from 6.55 – 14.27 leaves. The *CvLN* provides a measure of the genotype sensitivity to cool temperatures during the Veg phase. The *PvLN* and *PpLN* provide a measure of the photoperiod sensitivity of a genotype during the Veg and ER phases respectively. Separating responses during the Veg phase into cold and photoperiod responses and distinguishing photoperiod response between the Veg and ER phases is important. This is because photoperiod responses can work in opposite directions during these two phases and most other models are confounded because they fail to account for this. With the phenotyping scheme proposed here we demonstrate traits that related to gene expression controlling earliness *per se*, cold-temperature response (*Vrn1*), short-day vernalisation (*Vrn2*) and photoperiod sensitivity (*PpD*).

Alleles of the major developmental genes have been determined for each of the genotypes used in this study (Supplement SA1). However, there is substantially more variation in the phenotypes (Table 1) than can be related to these alleles alone. Other genes up- and downstream of the main control loop will also have an important role in the developmental phenotype. However, having a set of stable and unconfounded phenotypes provides a useful tool for genetic studies that aim to isolate and elucidate the role of different genes in wheat development.

**Table 1.** Allelic composition of wheat genotypes used in controlled environment phenology studies described in this paper.

|  | Genotype |  |  |  |  |  |
| --- | --- | --- | --- | --- | --- | --- |
| Marker | Otane | Batten spring | ‘Saracen’ | Batten winter | ‘Amarok’ | ‘CRWT15 3’ |
| *Vrn1* AF/R (Yan et al 2004) | A1a<br>(Spring Deletion) | A1a<br>(Spring Deletion) | A1<br>(Wild type – Winter) | A1<br>(Wild type – Winter) | A1<br>(Wild type – Winter) | A1<br>(Wild type – Winter) |
| *Vrn1* TDB Vrn-B1deletion* (Fu et al 2005) | Spring Deletion | Non Deletion | Non Deletion | Non Deletion | Non Deletion | Non Deletion |
| *Ppd-D1* F/R1/R2 (Beales et al 2007) | Insensitive | Sensitive | Sensitive | Sensitive | Insensitive | Sensitive |
| FT-A PJF3 allele seq (Bonnin et al 2008) | h4 | h1 | h3 | h1 | h3 | Not measured |
| FT-D PJF2 allele seq (Bonnin et al 2008) | h2 | h1 | h2 | h1 | h2 | Not measured |
| *Vrn2*ABD F1R2 (Zhu et al 2011) | ABD<br>(no null alleles) | ABD<br>(no null alleles) | ABD<br>(no null alleles) | ABD<br>(no null alleles) | ABD<br>(no null alleles) | ABD<br>(no null alleles) |

### 4.2 The link between *Vrn* gene expression and developmental phenotype

The CAMP model was developed to provide a quantitative framework linking genotype to developmental phenotype in a given environment through the expression of key development 25 genes. We used this model to explore the link between genotype (Table 1), observed developmental phenotypes (Figure 2 and Table 2), developmental physiology (Figure 3 and Figure 4) and foliar gene expression (Figure 5). The FLN-based phenotypes (Table 2) are an easily observable outcome of the timing of *VI^HS^* and *TS^HS^*, which are a less easily observable outcome of the expression of developmental genes. The gene expression depends on both the genetic makeup of a wheat plant and the temperature and photoperiod conditions it has grown through. The *FLN*-based phenotypes provide a relatively easy method of assessing the developmental phenotype which is important for large scale genomic studies. The results presented have shown the relationship between FLN and the effects of vernalization and photoperiod on *VI^HS^* and *TS^HS^*, assumed in the CAMP model, to be correct over a diverse range of genotypes (Figure 3). As such, we can focus the remainder of the discussion on the relationship between genotype, gene expression and the timing of *VI^HS^* and *TS^HS^*.

**Table 2.**
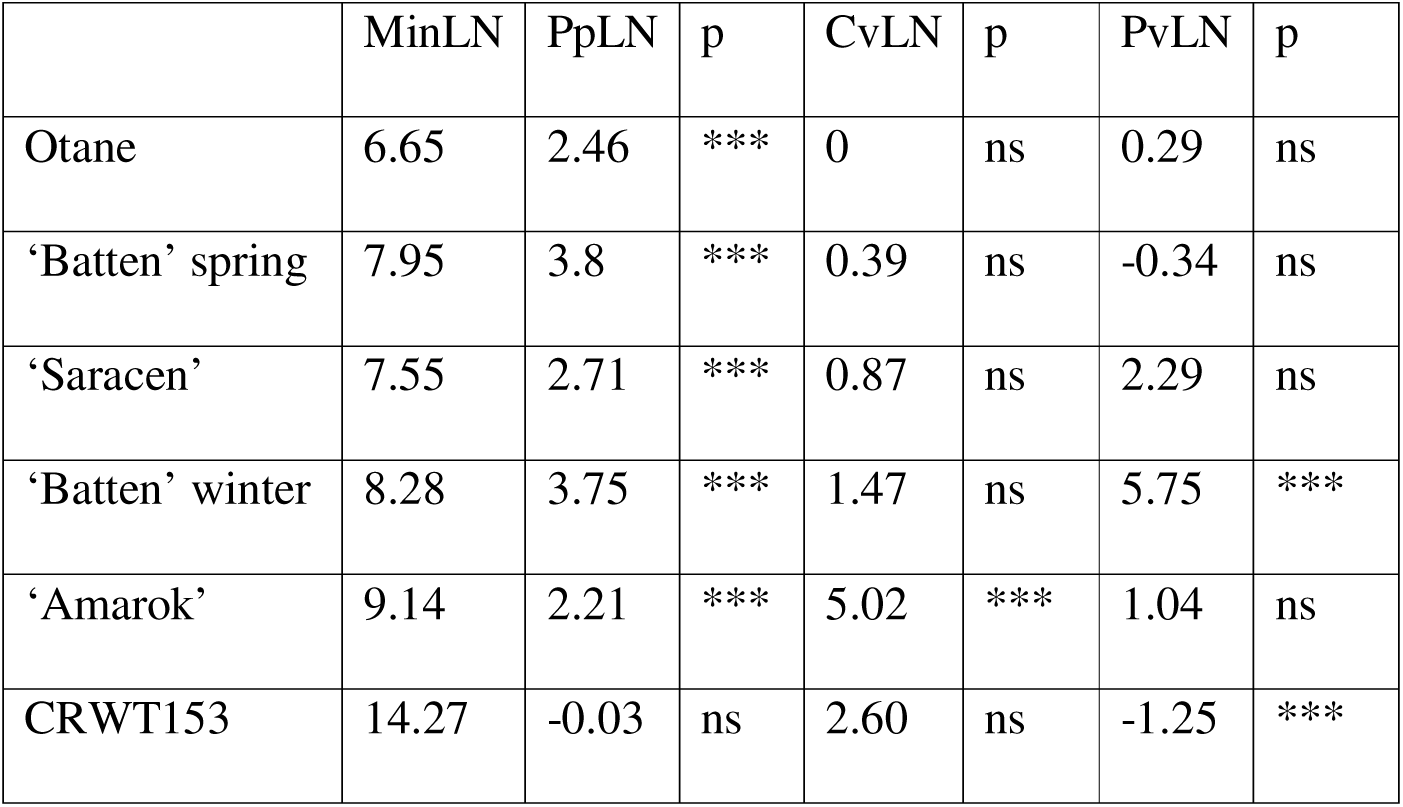
Developmental parameters for wheat genotypes derived from FLN observations conducted in controlled environment conditions. MinLN = minimum final leaf number; PpLN = photoperiod sensitivity; CvLN = cold vernalisation sensitivity; PvLN= photoperiod vernalisation sensitivity. Probability (p) levels are from t-tests of FLN data between the two treatments used to calculate the sensitivities (*** p <0.001, ** p <0.01, ns p > 0.05).

#### 4.2.1 *Vrn* gene expression and the timing of *VI^HS^*

The observed and predicted patterns of *Vrn* gene expression under CL conditions (Figure 5), are consistent with the molecular models of foliar interaction of these genes (Chen & Dubcovsky, 2012). Early cold treatments had a range of effectiveness for the winter genotypes shown in both the extent of early *Vrn2* and *Vrn1* expression (Figure 5) and the timing of *VI^HS^* (Table 1). The genetic control for this variation in response to cool temperature treatments is unknown, but it may relate to previously identified earliness *per se* loci (Sukumaran *et al*., 2016).

The lack of foliar *Vrn1* expression in the clear presence of *Vrn2* downregulation in the WL treatment is consistent with the suggestion of (Dubcovsky *et al*., 2006) of the existence of an alternative *Vrn2* repressor and with the results of Chen & Dubcovsky (2012) that foliar expression of *Vrn1* is not essential for reproductive transition. The patterns of foliar *Vrn* gene expression are consistent with the *Vrn-A1* and *Vrn-B1* alleles (Table 1). The lack of upregulation of foliar *Vrn1* in the winter genotypes suggests that the timing of *VI^HS^* is controlled by apical expression of *Vrn1* in combination with a *Vrn3* protein signal from the leaf. Upregulation of *Vrn3* was not observed in the winter genotypes in this treatment, nor was it observed in many of the S treatments, although it is considered to be an essential element of the transition to reproductive development (Chen & Dubcovsky, 2012). Our 26 results suggest that very low levels of foliar *Vrn3* expression (below the detection range of real time qPCR) are required to signal floral transition in the apex.

Foliar *Vrn1* was upregulated from the first leaf throughout development for the CS treatment but remained at relatively static levels because *Vrn3* was not upregulated (Figure 5). If we compare the CL treatment with the CS treatment, we see no difference in the time of *VI^HS^* (Figure 4) yet there are considerable differences in the patterns of foliar *Vrn2* and *Vrn3* expression (Figure 5). This further supports the suggestion that the timing of *VI^HS^* is controlled largely by *Vrn1* expression in the apex and that foliar expression of *Vrn* genes has little bearing, assuming a low expression of *Vrn3* occurs.

Finally, in the WS treatments (Figure 5), *Vrn1* is upregulated throughout for the spring genotypes, but only upregulated after *VI^HS^* or not upregulated at all for the winter genotypes. This further supports the notion that the timing of VS^HS^ is controlled largely by *Vrn1* expression in the apex and only a weak *Vrn3* signal is required from the leaf. In the WL treatments, we see *VI^HS^*occurring the latest, and genotypes where this is substantially later than under C treatments (i.e., the winter genotypes) all showed prolonged foliar expression of *Vrn2*. In all these cases, *Vrn2* was eventually downregulated which allowed low levels of *Vrn3* expression and the occurrence of *VI^HS^*.

Taken together, these results suggest that *Vrn1* expression in the apex is mostly responsible for triggering *VI^HS^* but foliar expression of *Vrn2* will keep *Vrn3* expression sufficiently low to prevent *VI^HS^* until it is all downregulated. When apical *Vrn1* is upregulated early, as in the C treatments and the spring genotypes under W, the pattern of *Vrn1* upregulation will be passed to the leaves that develop from the apex and this will hold *Vrn2* expression low and allow *VI^HS^* to occur. However, where *Vrn1* is not expressed early, some mechanism other than *Vrn1* repression of *Vrn2* is required to unlock the *Vrn3* signal from the leaves. Future improvements may be made to the CAMP model by explicitly modelling expression of *Vrn* genes in both foliar and apical sites and further investigating the quantitative link between absolute levels of gene expression and changes in apical morphological behaviour. This would enable more direct comparison between observed and modelled outcomes and further improvement of our understanding of the link between genetics and phenological development.

#### 4.2.2 *Vrn* gene expression, developmental phenotypes and the timing of *terminal spikelet*

Once the crop reaches *VI^HS^*, the role of foliar gene expression becomes clearer; there appears to be a clear link between the expression of foliar *Vrn3*, the duration from *VI^HS^* to *TS^HS^*, which is directly related to *PpLN* phenotype. CRWT153 had zero *PpLN* (Table 2) and this was explained because it did not appear to upregulate *Vrn3* either under long or short photoperiod conditions (Figure 5). The Batten genotypes had the greatest *PpLN* (Figure 2). This was because they did not express detectable levels of *Vrn3* under S conditions and there was a large extension of the duration from the *VI^HS^* to the *TS^HS^* phase (Figure 4). Otane and Amarok expressed some *Vrn3* under S conditions which would have caused some promotion of apical *Vrn1* (Yan *et al*., 2006), giving a smaller difference in *FLN* between long and short photoperiod treatments and a smaller *PpLN* (Table 2). The lower *PpLN* of Otane and Amarok corresponded with allelic variation in *Ppd-D1* (Table 1) but the genetic basis for the zero *PpLN* in CRWT153 is currently unknown.

The *CvLN* was calculated from the FLN difference between the CL and WL treatments (Figure 1). Comparing the *Vrn* expression patterns between these two treatments shows the spring types (Otane and Batten spring) upregulated *Vrn1* and subsequently downregulated *Vrn2* and upregulated *Vrn3* early regardless of vernalisation treatment. However, the winter types showed substantial and protracted foliar *Vrn2* expression and a lack of *Vrn1* and *Vrn3* expression under the WL treatment. The differences between winter and spring types related to differences in the *Vrn-A1* and *Vrn-B1* alleles (Table 1). There were substantial differences in the *VI^HS^* between ‘Saracen’ and the other winter types (Figure 4). This is related to low levels of foliar *Vrn1* expression for ‘Saracen’ compared with no foliar *Vrn1* expression for the other winter types. However, this variation in *CvLN* phenotype could not be related to the allelic makeup of these winter genotypes using published markers (Table 1) and further work is required to determine the genetic controls of this variation in vernalisation sensitivity.

The *PvLN* phenotype was relevant to the winter types (Batten winter, ‘Saracen’ and Amarok, Table 1) and its molecular cause can be clearly seen by comparing *Vrn* gene expression under WL and WS conditions (Figure 5). Batten winter showed the largest *PvLN* and under WL conditions (Figure 2) it showed substantial foliar *Vrn2* expression and no *Vrn1* or *Vrn3* expression. Under WS conditions Batten winter did not express foliar *Vrn2* and *VI^HS^*was substantially advanced (Figure 4). By contrast, Amarok had a small *PvLN,* expressed foliar *Vrn2* under both treatments and showed a smaller reduction in *VI^HS^*. The difference in *Vrn2* expression between Batten winter and Amarok was probably caused by differences in the *Ppd-D1* allele (Table 1) which sits upstream of *Vrn2* expression (Mulki & von Korff, 2016). This shows another advantage of using the controlled environment phenotyping method described here, where the confounding effects of photoperiod and vernalisation responses were clearly separated. The non-vernalised winter genotypes did not upregulate *Vrn1* under either short or long photoperiod after *Vrn2* was downregulated, which is consistent with the results of (Dubcovsky *et al*., 2006).

### 4.3 Concluding remarks

The data presented by Brooking & Jamieson, (2002) and the analyses conducted by Brown *et al*., (2013) on the environmental responses of, and relationships between *FLN*, *TS^HS^* and *VI^HS^*for the Batten near isogenic pair were replicated in the current work. We modified the original CAMP conceptual model (Brown *et al*., 2013) to better capture how expressions of *Vrn1*, *Vrn2* and *Vrn3* influence development progress. The analysis presented here showed that the responses and relationships hold for a broad range of phenotypes demonstrating the versatility of the phenological framework in the modified CAMP model. Comparison of the relationships between environmental conditions, *Vrn* gene expression and the timing of key stages of apical development are consistent with those predicted by the new CAMP model. This supports its use to provide stable phenotypes for characterising wheat genotypes. Specifically, exposure to cool temperature at early stages of development (before HS – 2.0) upregulated *Vrn1* and accelerated the onset of *VI* in winter wheat types. *Vrn1* was upregulated and *VI* occurred early in spring wheat types regardless of the temperature conditions encountered. *Vrn*2 and *Vrn3* were only upregulated under long photoperiod for most cultivars. The upregulation of *Vrn1* was accompanied by a downregulation of *Vrn2*. We propose that *FLN*-based phenotypes, assessed with a factorial of plus and minus early cold treatment and long and short photoperiod, provide unconfounded phenotypes that will be useful in large scale genomic studies. The CAMP model also provided a useful framework for reconciling the complex linkage between genotype, gene expression, apical physiology and the resulting G×E patterns observed in *FLN* in wheat. The APSIM wheat model (Brown *et al*., 2018) has been upgraded to include the CAMP phenology model and a full description of this process is given in an accompanying paper (Wang *et al*., 2025)

## 5. Acknowledgments

Experimental and conceptual model development was completed under The New Zealand Institute for Plant and Food Research Limited Sustainable Agro-ecosystems (SAE) programme, with funding from the Ministry of Business, Innovation and Employment (MBIE) Strategic Science Investment Fund (SSIF). The SAE programme focussed on delivering transdisciplinary scientific knowledge, tools and technologies that enhance the productivity and resilience of primary industries while reducing their environmental footprint to meet community and market defined limits.

Refinement and completion of the CAMP model was made possible by investment from the Grains Research and Development Corporation (GRDC) through the National Phenology Initiative (ULA00011), and the field support from the South Australian Development and Research Institute agronomy team.

## 7. Author contributions

HB was the initiator of the CAMP model concept and the experimental research presented in this paper, designed, implemented and monitored the experimental work, analysed the data, wrote prototype model code, produced all the graphics produced the first paper draft. EW, JM, DM, RM and NH provided input to the interpretation of results and improvement of the model. PJ provided genetic material and allelic composition data. MPJ provided guidance on tissue sampling and completed qPCR extraction and analysis. BZ and ZZ provided implementation and testing of the model in broader contexts. EW contributed substantially to the improvement of model concepts and the manuscript and all authors provided final checking.

## 8. Data Availability

All the data and scripts used to analyse data and produce graphs as well as CAMP model code are publicly available at https://github.com/HamishBrownPFR/CAMP/

