## Supplement for "Integrating molecular and physiological approaches to quantify genetic controls for wheat development and improve phenotyping"

### **Supplementary Materials**

A. PCR primers used in this study
B. Observations of diurnal gene expression patterns
C. Description of the Cereal Anthesis Molecular Phenology (CAMP) models algorithms and parameterization schemes.

Supplement A: PCR primers used in this study.

Table SA1. Description of primers used.

| **Gene** | **Genbank Accession** | **Annotation** | **5′–3′ Sequence** |
| --- | --- | --- | --- |
| *Vrn1* | AB007504 | MADS box transcription factor | Forward GGAGAGGTCACTGCAGGAGGA  Reverse GCCGCTGGATGAATGCTG |
| *Vrn2* | AY485965 | Zinc finger-CCT domain gene | Forward AGCCCACATCGTGCCATTT  Reverse ACCTCATCACCTTCGCTGCT |
| *Vrn3* | AB361063 | Triticum aestivum WFT mRNA for putative kinase inhibitor | Forward CGCTGGTGGTTGGCAGGGTT  Reverse TCACAAGCCAGTGGAGATACTCCCTAA |
| *Ta54227* | Ta54227 | Cell division control protein | Forward CAAATACGCCATCAGGGAGAACATC  Reverse CGCTGCCGAAACCACGAGAC |
| *Tef-1α* | M90077 | Translation elongation Factor 1 alpha-subunit (TEF1) | Forward CTGCTGCAACAAGATGGACG  Reverse GAGATGGGGACGAAGGGAAC |

Supplement B: Diurnal patterns of Vrn gene expression.

A preliminary experiment was conducted to aid the protocol development for sampling foliar expression of Vrn genes. The diurnal patterns of gene expression of Otane, Batten spring, ‘Saracen’ and Batten winter genotypes were measured on two occasions at *HS*=3 and *HS*=9 at 23°C under 16 h photoperiod. Leaf tissue was harvested at 2-hourly intervals over two 24-hour periods 16, 46 days after planting (representing Haun leaf stage of 3 and 9 respectively). For RNA isolation, the most recent fully expanded leaf was cut (using sharp scissors) at its mid length and immediately, ~5 × 1-mm strips ( ~1 cm^2^,  ~25 mg) were cut from the basal end of the sampled leaf directly into a 1-mL tube containing 300 µL of DNA/RNA Shield (Zymo Research Ltd.). Care was taken to completely submerge the leaf tissue in the DNA/RNA shield. Samples were stored at −20^°^C for later RNA isolation

Overall patterns of *Vrn* expression at *HS*=3 were consistent (Figure SB1) with the earliness of varieties (See main body of paper); Otane was earliest (*FLN* = ~7) and would have passed *VI^HS^* by this point. It had the highest *Vrn1* and *Vrn3* expression and minimal *Vrn2*. Batten spring was next earliest (*FLN* = ~8) with the next highest *Vrn1* and *Vrn3* expression, ‘Saracen’ (*FLN* = ~10) and Batten winter (*FLN* = ~16) were the latest varieties and showed the highest *Vrn2* and lowest *Vrn1* expression at *HS* = 3. There was a clear diurnal pattern showing a gradual decrease in *Vrn1* from the start to the end of the day suggesting there is a rapid upregulation of foliar *Vrn1* at the start of the light period. ‘Saracen’ showed a different pattern of *Vrn2* expression compared with the other genotypes with a substantial up-regulation later in the light period.

At *HS*=9 both Otane and Batten spring were at flag leaf and ‘Saracen’ was past terminal spikelet so foliar gene expression profiles had little relevance to the reproductive development of the plant. However, diurnal patterns of gene expression were similar to those at *HS*=3. The relative expression of a second house-keeper gene (*Tef-1α*) is also shown to provide a comparison of the temporal stability of the two house-keeper genes (Figure SB1). While there were no genotype differences apparent in *Tef-1α* expression, there were considerable diurnal variations in relative expression. Overall, the expression at any time of day correlates well with the expression integrated over the entire day, suggesting representative samples could be collected at any time during the light period. We settled on the end of the light period for our sampling in the main experiment.

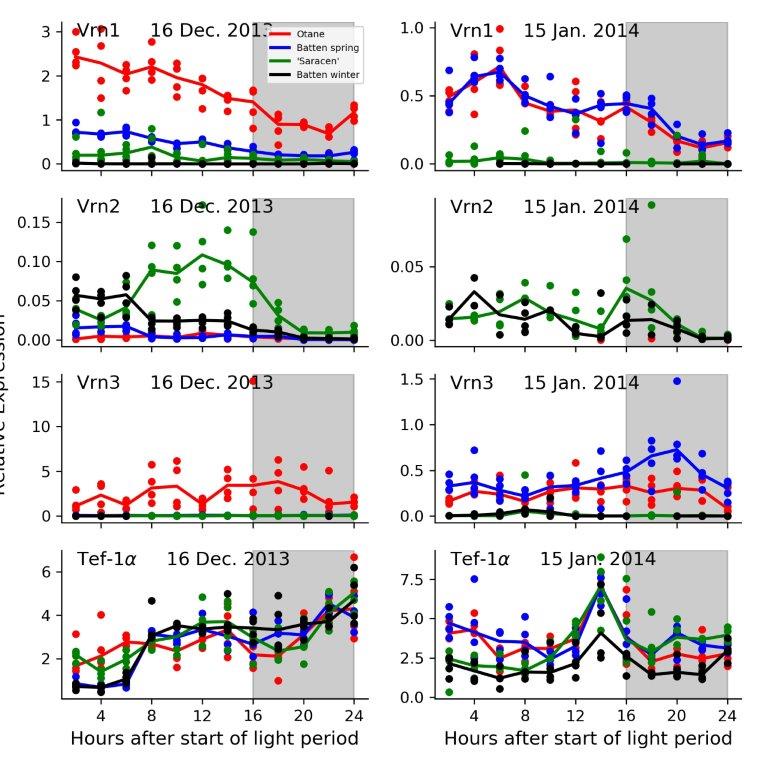

Figure SB1. Diurnal patterns of gene expression (relative to the house-keeping gene, *Ta54227*) for different genotypes measured on two dates when the plants had 3 (16 Dec. 2013) and 9 (15 Jan. 2014) main-stem leaf ligules. Grey shading represents the dark period.

### Supplement C: Description of the Cereal Anthesis Molecular Phenology (CAMP) models algorithms and parameterization schemes.

#### 1.1 Description of the CAMP model

The CAMP (Cereal Anthesis Molecular Phenology) model integrates physiological and molecular models of anthesis to disentangle the G x E mediated processes that underly the timing of anthesis. It draws on the hypothetical model of Brown *et al.*, (2013), with substantial changes in the conceptualization and implementation. Firstly, the thermal time accumulated between sowing and anthesis (*Sow_Anthesis^tT^*) can be calculated as the sum of thermal time between sowing and flag leaf expanded (*Sow_FlagLeaf^Tt^*) and that between flag leaf expanded and anthesis (*FlagLeaf_Anthesis^Tt^*):

*Equation 1*	${Sow\_Anthesis}^{Tt}={Sow\_FlagLeaf}^{Tt}+FlagLeaf\_{Anthesis}^{Tt}$

Where sow_FlagLeaf^Tt^ is calculated as:

Equation 2	$Sow\_{FlagLeaf}^{Tt}=FLN \times Phyllochron$

Where Phyllochron is the thermal time interval between the appearance of two consecutive leaves on the main stem, representing a genotype specific rate of leaf appearance in response to temperature accumulation and:

Equation 3	$FLN=2.85+ {1.1\times TS}^{HS}$

TS^HS^ is assumed to occur when an apical development index (*ApDev_d_*) reaches a value of 2.0. *ApDev_d_* is calculated each day until TS^HS^ as:

Equation 4	${ApDev}_{d}= min({MaxVrn}_{d}, {VrnB}_{d}+ {Vrn1}_{d}+{Vrn3}_{d}-{MaxVrn2}_{d})$

Where ${MaxVrn}_{d}$ is the maximum daily developmental progress possible in response to Vrn gene expression (Equation 5) which represents a morphological limit to the speed at which the plant can physically exhibit responses to gene expression. *VrnB_d_* is the base expression of protagonistic development genes that occurs in the absence of any cold-induced upregulation of Vrn1 or photoperiod induced upregulation of Vrn3 (Equation 6). *Vrn1_d_* is the persistent cold-induced upregulation of the cold vernalization signal (Equation 8). Please note that in the associated paper we treat VrnB and Vrn1 as a single entity but separate them in the models algorithms for numerical clarity. *Vrn3_d_* is photoperiod induced upregulation of the Vrn1 promoter (Equation 13-14). Max*Vrn2* is the antagonistic effect of Vrn2 expression (Equation 10).

Equation 5	${MaxVrn}_{d}={MaxVrn}_{d-1}+rVrnM\times{\Delta HS}_{d}$

Where *rVrnM* is a cultivar specific coefficient quantifying the maximum potential development rate and may have different values before and after VI. ΔHS_d_ is the daily progression of Haun Stage.

The period of crop development preceding TS^HS^ is subdivided into three phases;

- Emergence phase (Imbibition to Emergence) when *VrnB_d_* and *Vrn1_d_* upregulation may occur.
- Vegetative phase (Emergence to VI^HS^) when *VrnB_d_, Vrn1_d_, Vrn2_d_* and *Vrn3_d_* upregulation may occur.
- Early Reproductive Phase (VI^HS^ to TS^HS^) when *VrnB_d_* and *Vrn3_d_* upregulation may occur.

We assume VI^HS^ and TS^HS^ occur at arbitrary *ApDev_d_* thresholds of 1.0 and 2.0, respectively. *VrnB_d_* will accumulate each day through all three phases and is calculated as:

Equation 6	${VrnB}_{d}={VrnB}_{d-1}+{\Delta VrnB}_{d}$

Equation 7	${\Delta VrnB}_{d} = rVrnB \times{\Delta HS}_{d}$

rVrnB is the VrnB expression rate coefficient for the given genotype and may have different values before and after VI^HS^. Accumulated persistent Vrn1 expression is calculated in response to accumulated cold perception (*cold_d_*) as:

Equation 8	${Vrn1}_{d}=max(0,{(cold}_{d-1}+ {\Delta cold}_{d})-MT)$

Where MT is a methylation threshold for how much effective cold must be perceived before Vrn1 expression is methylated and cold response becomes persistent. Following Brown *et al.*, 2013) MT has a value of half of the accumulated *cold_d_* required to saturate cold-temperature responses. The daily increment of cold perception (Δcold_d_) is calculated as:

Equation 9	${\Delta cold}_{d}=\exp\left( -0.18 \times T \right)\times rVrn1\times{\Delta VrnB}_{d}$

Where T is temperature, and rVrn1 is a genotype specific coefficient that quantifies the upregulation of Vrn1, relative to *VrnB_d_* expression, by exposure to cold.

It is assumed that *Vrn2_d_* is rapidly upregulated soon after the perception of long photoperiod to a level that is subsequently up- or downregulated dependent on photoperiod:

Equation 10	${MaxVrn2}_{d}=max(0,mVrn2\times{fPP}_{d}-{AccumSPp}_{d})$

Where *mVrn2* is the genotype-specific coefficient quantifying the maximum Vrn2 expression under long days and *fPP* is calculated from photoperiod, increasing linearly from 0.0 at Pp≤8 h to 1.0 at Pp≥16 h. *AccumSPp* reduces MaxVrn2 if long photoperiod is perceived after a period of short photoperiod:

Equation 11	${AccumSPp}_{d}= {AccumSPp}_{d-1}+ {\Delta VrnB}_{d} \times(1-fPP)$

This is an empirical function to capture the reduced effectiveness of long days to delay VI^HS^ following an initial period of exposure to short days (Brooking & Jamieson, 2002) and further investigation is required to understand the molecular basis for this response.

Vrn2 is blocked by base and cold Vrn expression so its apparent expression on any day (d) is calculated as:

Equation 12	${Vrn2}_{d}=max({MaxVrn2}_{d}-\left( {VrnB}_{d}+{Vrn1}_{d} \right),0)$

Vrn3 is expressed when ${Vrn2}_{d}$ = 0 and is calculated as:

Equation 13	${Vrn3}_{d}={Vrn3}_{d-1}+ {\Delta Vrn3}_{d}$

Where ΔVrn3_d_ is calculated as:

Equation 14	${\Delta Vrn3}_{d}=(rVrn3-1)\times fPP\times{\Delta VrnB}_{d}$

Where rVrn3 is a genotype-specific coefficient that describes the increase in Vrn3 expression, relative to VrnB expression, with increasing photoperiod and may have different values before and after VI^HS^.

Equations 4 – 14 are solved each day until *ApDev_d_* ≥ 1.0 (i.e. VI^HS^ is reached) then *ApDev_d_* is calculated as:

Equation 15	${ApDev}_{d}={ApDev}_{d-1}+{\Delta VrnB}_{d}+{\Delta Vrn3}_{d}$

Equation 15 is solved until *ApDev_d_* is ≥ 2.0 at which time TS^HS^ is known, FLN can set (Equation 3) and *Sow_FlagLeaf^Tt^* can be calculated using Equation 2.

Finally *FlagLeaf_Anthesis^Tt^* (Equation 1) is calculated from a base thermal time duration that occurs under long photoperiod (*LongPpBase*) and any additional thermal time that results from shorter photoperiod (*ShortPpExtension*). Both the *LongPpBase* and the photoperiod sensitivity parameter in calculating *ShortPpExtension* are genotype specific cultivars

#### 1.2 Deriving genotype specific coefficients for CAMP

The CAMP gene expression model (Equations 4-15) requires 9 genotype specific coefficients:

1) ${rVrnB}_{Veg}$: base expression of main development genes in vegetative stage
2) ${rVrnB}_{ER}$: base expression of main development genes in early reproductive stage
3) ${rVrnM}_{Veg}$: maximum daily developmental progress in vegetative stage
4) ${rVrnM}_{ER}$: maximum daily developmental progress in early reproductive stage
5) $rVrn1$: upregulation of Vrn1 by cold, relative to *VrnB_d_*
6) $MT:methylation threshold of percieved cold$
7) $mVrn2$: maximum Vrn2 expression under long days
8) ${rVrn3}_{Veg}$: upregulation of Vrn3 by photoperiod in vegetative stage, relative to VrnB expression
9) ${rVrn3}_{ER}$: upregulation of Vrn3 by photoperiod in early reproductive stage, relative to VrnB expression

These can be calculated from genotype specific measures of phyllochron and FLN measured in the four different environments (E) where plants are exposed to cool (C) or warm (W) temperatures between germination and VI^HS^ in combination with short (S) or long (L) photoperiods from emergence until flag leaf expended. These treatments are represented by the symbols CS, CL, WS and WL.

##### 1.2.1 Deriving VI and TS timings from FLN observations

First of all TS^HS^ is calculated for each environment:

Equation 16.	${{TS}^{HS}}_{E}={(FLN}_{E}-2.85)/1.1$

Which is Equation 3 rearranged to make TS^HS^ the subject.

Then the timing of VI^HS^_E_ is calculated from TS^HS^_E_ for each environment using the following calculation scheme. This scheme is also drawn in Figure SC1. For the CL and WL treatments we assume the duration in Haun stage (HS) of the early reproductive phase (ER^HS^ = TS^HS^- VI^HS^) will be minimized (MinER^HS^) because long photoperiod is maximizing expression rates. For the CS and WS treatments we assume ER^HS^ will be maximized (MaxER^HS^) as expression rates will be minimized under short photoperiod. Thus, VI^HS^_E_ are calculated as follows. Firstly, MinER^HS^ is calculated as:

Equation 17.	$\mathrm{MinER}^{HS}= min(3,{\mathrm{TS}^{HS}}_{CL}-\mathrm{MinVI}^{HS})$

Where upper bound of 3 represents the longest duration ER^HS^ recorded in long photoperiod treatments. MinVI^HS^ is the soonest a wheat plant can display VI^HS^ and is represented by a value of 1.1 which was the smallest value observed.

Next, we subtract MinER^HS^ from the TS^HS^_E_ for the long photoperiod treatments to estimate VI^HS^:

Equation 18.	${{VI}^{HS}}_{CL}= {{TS}^{HS}}_{CL}-\mathrm{MinER}^{HS}$

Equation 19.	${{VI}^{HS}}_{WL}=\max\left( {{TS}^{HS}}_{WL}-\mathrm{MinER}^{HS},{{VI}^{HS}}_{CL} \right)$

The max constraint is applied to VI^HS^_WL_ to ensure it is not earlier than VI^HS^_CL_ in the few cases where FLN_WL_<FLN_CL_. Vrn2 is inactive under short photoperiod conditions so VI^HS^ will not be delayed and we assume that VI^HS^_CS_ will be at least as early as VI^HS^_CL_. It is possible cold vernalization was incomplete at the end of the cool temperature treatment and some Vrn2 expression occurred prior to VI^HS^_CL_. If this were the case, VI^HS^_CS_ would be less than VI^HS^_CL_. However, the situations where VI^HS^ was observed from destructive sampling (Brown *et al.*, submitted) showed it was the same under both CL and CS treatments. This was even true for CRWT153 which had VI^HS^ of 9.0 after cool vernalization treatments. So we assume Vrn2 expression is negligible in the CL treatment and the MinER^HS^ calculated in Equation 17 is also applicable for estimating VI^HS^_CS_:

Equation 20.	${{VI}^{HS}}_{CS}= min( {{TS}^{HS}}_{CS}-\mathrm{MinER}^{HS},{{VI}^{HS}}_{CL})$

The min constraint is included for the few cases where FLN_CS_ < FLN_CL_. We can now calculate MaxER^HS^ as:

Equation 21.	$\mathrm{MaxER}^{HS}= {\mathrm{TS}^{HS}}_{CS}-{\mathrm{VI}^{HS}}_{CS}$

Assuming the ER^HS^ will be the same in both short photoperiod treatments, we can calculate VI^HS^_WS_ as:

Equation 22.	${{VI}^{HS}}_{WS}= max({{TS}^{HS}}_{WS}-\mathrm{MaxER}^{HS},{{VI}^{HS}}_{CL})$

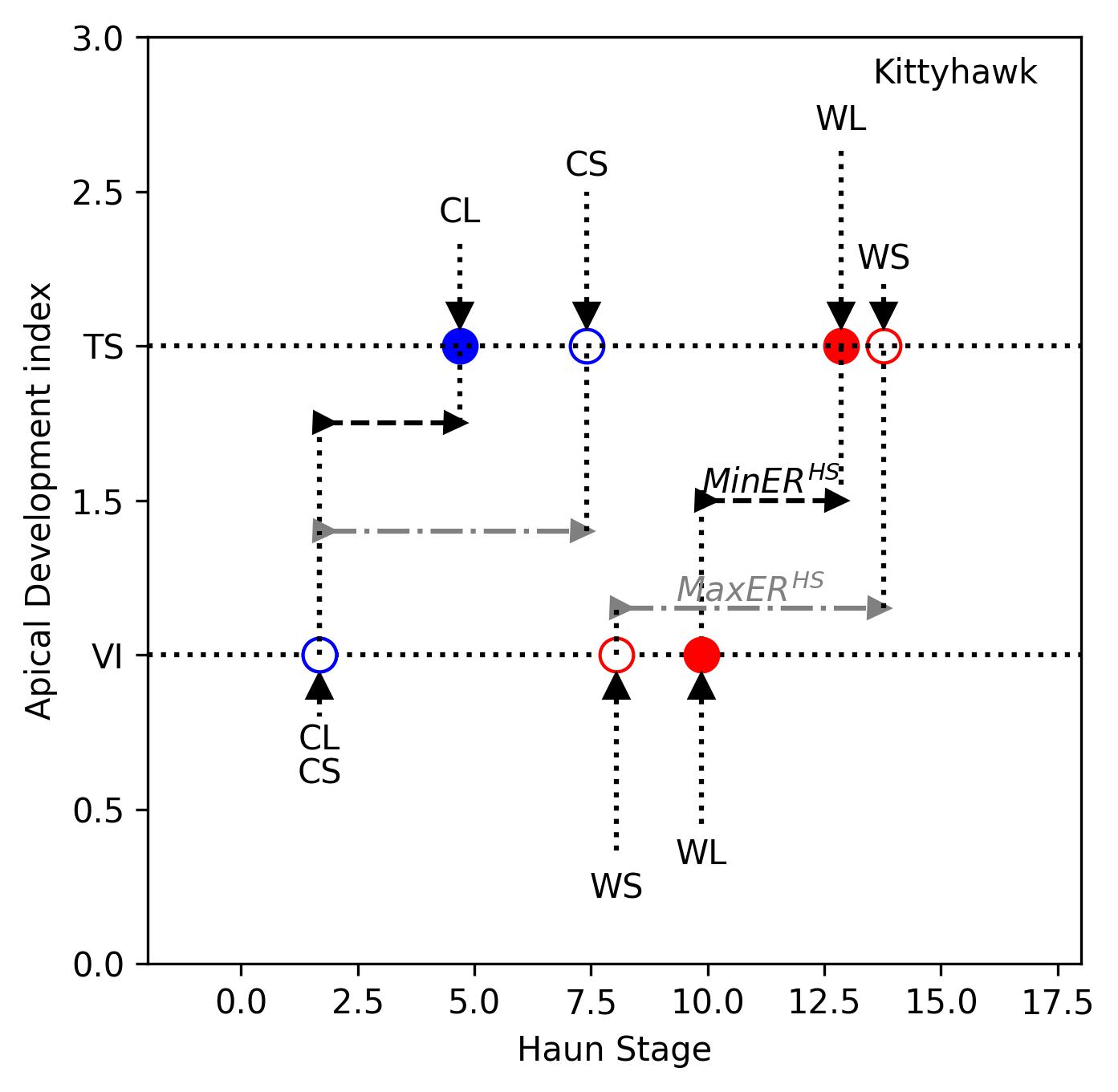

Figure SC1. Plot of the coordinates of vernal induction (VI) and terminal spikelet (TS) for wheat cultivar Kittyhawk (Bloomfield *et al.*, 2023) under the four treatments (CL – cold & long photoperiod (Pp), CS – cold & short Pp, WS – warm & long Pp, and WL – warm & long Pp). The Haun stage duration of the early reproductive phase (*ER^HS^ = TS^HS^- VI^HS^*) is minimized under CL and WL (*MinER^HS^*), while maximized under CS and MS (*MaxER^HS^*).

##### 1.2.2 Deriving base and maximum expression rates

Now we have established VI^HS^ and TS^HS^ for each of the four environments, and if we assume that VI^HS^ occurs at an apparent Vrn expression (*ApDev_d_*) of 1.0 and TS^HS^ occurs at *ApDev_d_* of 2.0, we can have coordinates for calculating the base and maximum bounds for apparent expression (Figure SC2). In the WS treatment there is no cold acceleration of Vrn and no Vrn2 or Vrn3 upregulation to influence development so Vrn expression will be at a base rate calculated for the vegetative phase (*rVrnB_veg_*) as:

Equation 23.	*rVrnB_Veg_* = 1 / (*VI^HS^_WS_ + ED^HS^*)

Where *ED^HS^* is the Haun stage equivalent duration of emergence calculated as the thermal time from germination to emergence divide by the phyllochron. The rate for the early reproductive phase (*rVrnB_ER_*) is calculated as:

Equation 24.	*rVrnB_ER_* = 1 / (*TS^HS^_WS_* - *VI^HS^_WS_*)

The CS treatment has Vrn1 upregulated to its maximum and no upregulation of Vrn2. Thus, VI^HS^_CS_ occurs at the earliest possible HS and, we can use this treatment to quantify the maximum apparent Vrn expression during the vegetative phase (*rVrnM_Veg_*):

Equation 25.	*rVrnM_Veg_* = 1 / (*VI^HS^_CS_* + *ED^HS^*)

During early reproductive phase, development will progress at its maximum under long Pp where Vrn3 is upregulated. Thus, maximum apparent expression during this phase (*rVrnM_ER_*) is calculated as:

Equation 26.	*rVrnM_ER_* = 1 / *MinER^HS^*

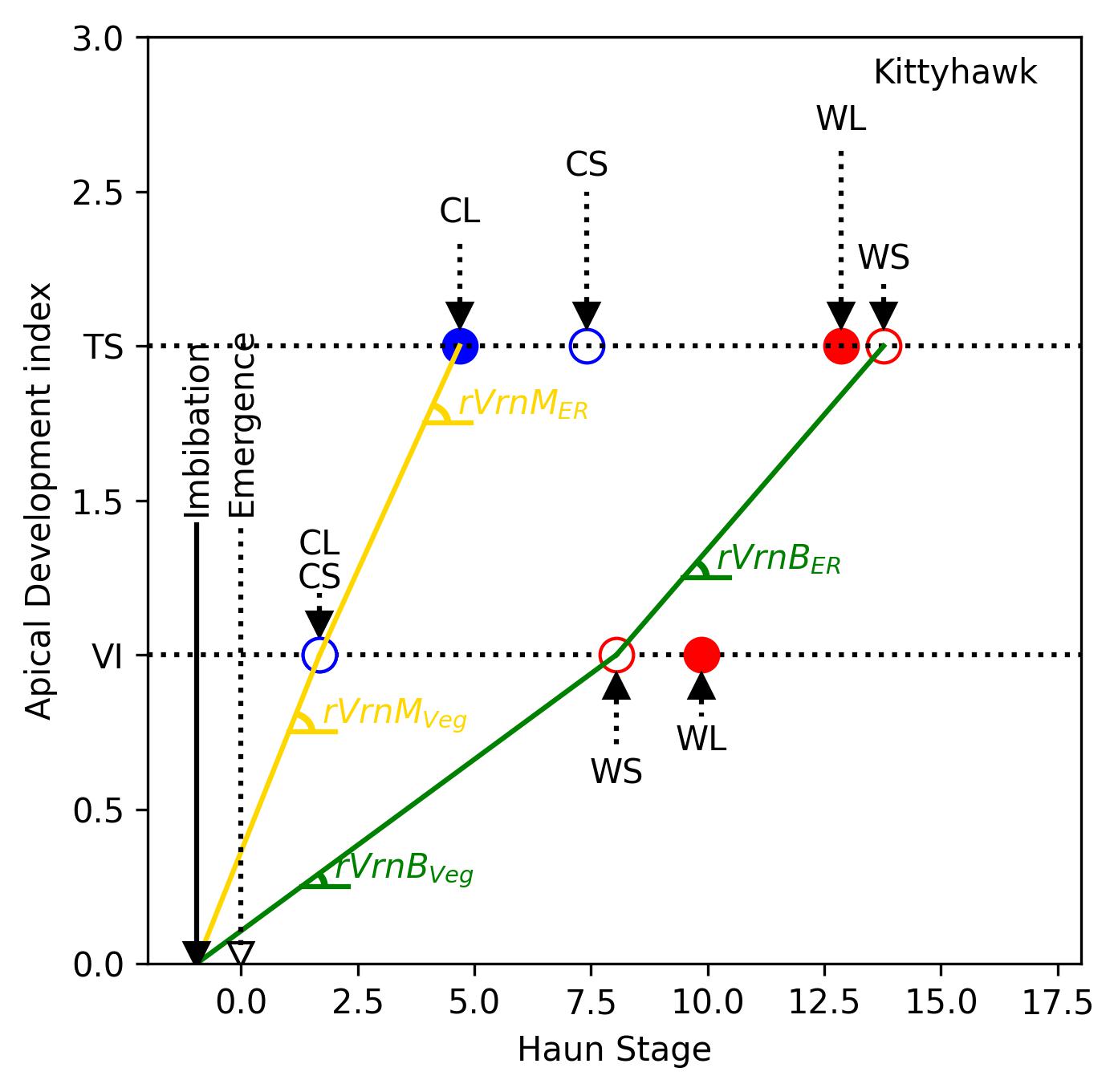

Figure SC2. Plot of vernal induction (VI) and terminal spikelet (TS) for wheat cultivar Kittyhawk (Bloomfield *et al.*, 2023) against Vrn expression under the four treatments (CL – cold & long photoperiod (Pp), CS – cold & short Pp, WS – warm & long Pp, and WL – warm & long Pp). The base rates were derived from WS treatment, and the maximum rate of expression was derived from CS treatment for vegetative stage and from the CL treatment for the early reproductive stage.

##### 1.2.3 Deriving Vrn3 expression coefficients

Now that we have established the base and maximum boundaries to apparent Vrn expression, we can derive the parameters that describe the effects of environment on expression of Vrn1, Vrn2 and Vrn3 that modulate expression within these bounds.

Firstly, we derive factors for the photoperiod sensitivity of Vrn3 expression in the early reproductive phases (*rVrn3_ER_*):

Equation 27.	*rVrn3_ER_* = *rVrnM* *_ER_* / *rVrnB_ER_*

Such that a value of 1.0 will mean there is no Vrn3 expressed above base rate (i.e. the genotype is photoperiod insensitive) and a value > 1.0 shows increasing upregulation of Vrn3 by long photoperiod.

We do not have sufficient information in the coordinate scheme (Figure SC2) to derive Vrn3 sensitivity in the vegetative (rVrn3_Veg_) phase explicitly. Instead, we assume the rVrn3_Veg_ is the same as that derived for the ER phase (Equation 27). The only exception to this is for genotypes that have an early VI^HS^_WL_ that requires a rate of Vrn3 expression greater than that given by rVrn3_ER_. To capture this rVrn3_Veg_ is calculated as the maximum of rVrn3_ER_ and the rate of Vrn3 expression that is needed to occur to achieve VI^HS^_WL_:

Equation 28.	*rVrn3_Veg_* = max(*rVrn3_ER_*,(1 / *VI^HS^_WL_*) / *rVrnB_Veg_* )

##### 1.2.4 Deriving Vrn2 expression rates

The next parameter required is the maximum Vrn2 expression under long photoperiods (*mVrn2*). A visual representation of the scheme used to calculate this is given in (Figure SC3). In the WL treatment, exposure to long photoperiod following emergence will firstly delay progress toward VI through expression of Vrn2 which blocks Vrn3. When Vrn2 is effectively downregulated, Vrn3 will be upregulated by long photoperiod which will accelerate progress toward VI. The HS duration of the period of Vrn3 upregulation prior to VI (${{URVrn3}^{HS}}_{WL}$) is given as the amount of Vrn upregulation (VI target = 1.0) over the rate of expression:

Equation 29.	${{URVrn3}^{HS}}_{WL}=1/\left( r{VrnB}_{Veg} {\times rVrn3}_{Veg} \right)$

If we subtract this duration from VI^HS^_WL_, we will get the duration of the period of Vrn2 downregulation (*DRVrn2^HS^_WL_*):

Equation 30.	*DRVrn2^HS^_WL_* = Max(0, *VI^HS^_WL_* – *URVrn3^HS^_WL_*);

Then we assume that Vrn2 is downregulated at a rate equivalent to ΔVrnB_veg_ (Equation 7) under WL (at the base rate because of no vernalization under WL), we can extrapolate ΔVrnB_veg_ expression back over DRVrn2^HS^_WL_ at this rate to determine what the expression of Vrn2 would have been near emergence (mVrn2).

Equation 31.	*mVrn2* = (*DRVrn2^HS^_LN_* + *ED^HS^*) $\times$ *rVrnB_Veg_*.

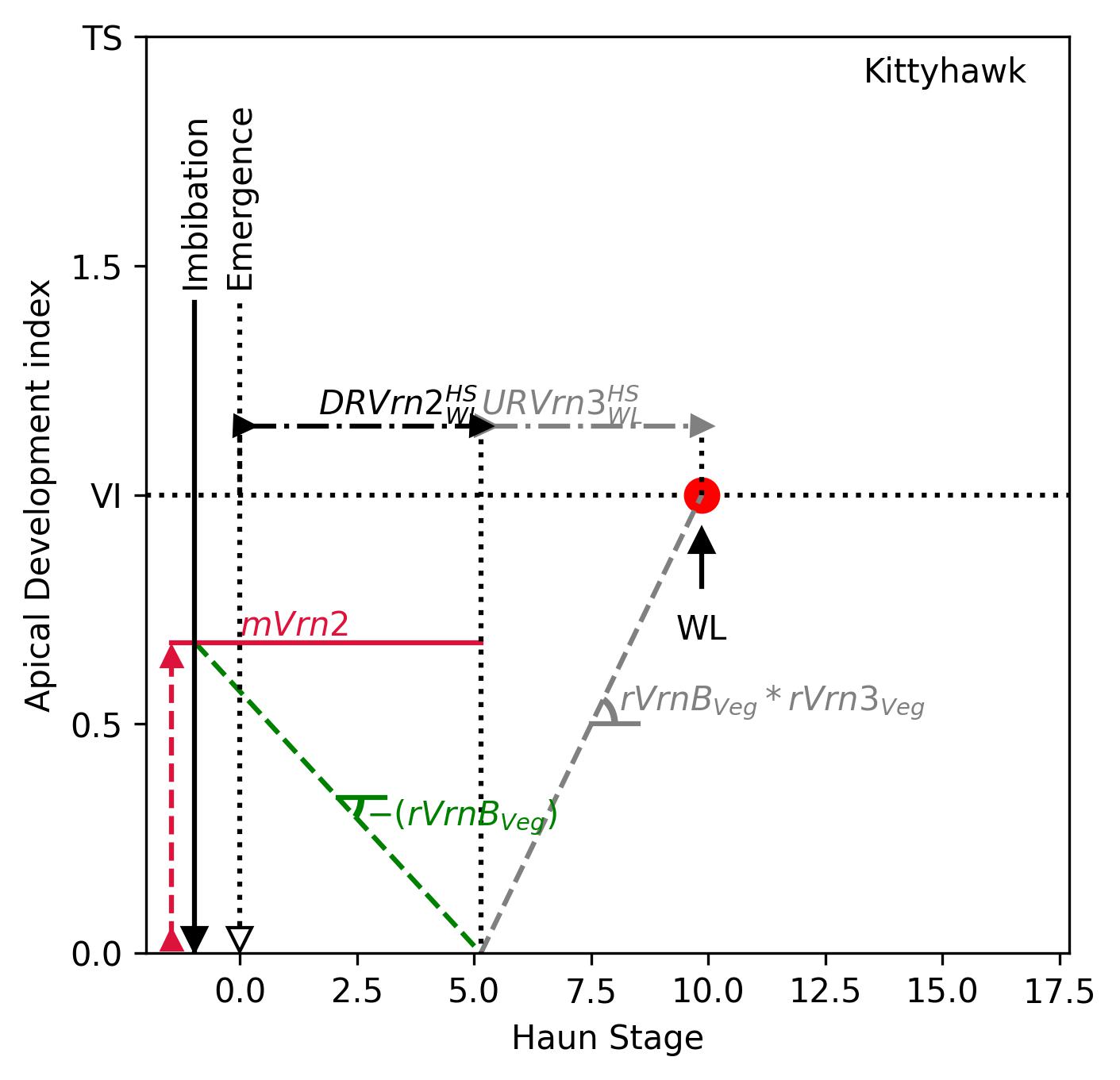

Figure SC3. Plot demonstrating calculation of maximum Vrn2 expression (*mVrn2*) under treatment WL for wheat cultivar Kittyhawk (Bloomfield *et al.*, 2023). First, extrapolating Vrn3 expression back from VI^HS^_WL_ to determine the duration when Vrn3 is upregulated by photoperiod prior to VI (*URVrn3^HS^_WL_*), then by difference calculating the duration for Vrn2 to be down regulated (*DRVRN2^HS^_WL_*). Assuming Vrn2 is downregulated under the base Vrn rate (due to no vernalizing temperature under WL), then extrapolating base Vrn expression up enables estimation of Vrn2 expression levels at emergence (*mVrn2*). More details see text.

##### 1.2.5 Deriving Vrn1 expression coefficients.

The final parameter we need to derive quantifies the cold-vernalization sensitivity of the genotype and the scheme to calculate this is shown graphically in Figure SC4. We assume the protagonistic Vrn expression required to achieve VI under long day conditions (*Vrn^P^VI_CL_*) is the VI threshold (1.0) and the amount of *Vrn^P^* needed to offset antagonistic Vrn2 expression (*mVrn2* under long day conditions):

Equation 32.	*Vrn^P^VI_CL_* = 1.0 + *mVrn2*.

We assume Vrn3 expression will be negligible prior to VI in the CL treatment so protagonist gene expression will be from VrnB and Vrn1. Therefore, the amount of persistent Vrn1 expression (${pVrn1}_{\mathrm{CL}}$) required is calculated as:

Equation 33.	$p{Vrn1}_{CL}= {Vrn}^{P}{VI}_{CL}-\sum_{VI}^{0} {VrnB}_{Veg}$

Where $\sum_{VI}^{0} {VrnB}_{Veg}$ is the amount of base vernalization expressed from imbibition to VI and is calculated as:

Equation 34.	$\sum_{VI}^{0} {VrnB}_{Veg} = ({ED}^{HS} + {{VI}^{HS}}_{CL}) \times{rVrnB}_{Veg}$

To achieve persistent (methylated) Vrn1 expression we assume unmethylated Vrn1 must first be upregulated to a methylation threshold (MT) and any subsequent Vrn1 expression is methylated. This produces the vernalization lag we see in experiments where short vernalization treatments give no reduction in final leaf number.

Based on the data of Brooking and Jamieson (2002) the lag duration is the same length as the duration from the end of the lag until cold vernalization response is completed. Therefore, we assume the MT is the same as the amount of Vrn1 that is required to give full persistent vernalization response:

Equation 35.	*MT* = $p{Vrn1}_{CL}$

The duration over which Vrn1 upregulation occurred (Vrn1^HS^) is the minimum of the vernalization treatment duration (VD^HS^) and the sum of VI^HS^_CL_ and ED^HS^ (for cases where VI occurred before the end of the vernalization treatment) calculated as:

Equation 36.	*URVrn1^HS^_CL_* = min(*VD^HS^*, *VI^HS^_CL_* + *ED^HS^*);

Then we can calculate the rate of Vrn1 expression in response to cool treatment temperature (ΔVrn1_CL_) as:

Equation 37.	*ΔVrn1_CL_* = ($p{Vrn1}_{CL}$+ *MT*) / *URVrn1^HS^_CL_*

We assume an exponential decline in Vrn1 expression with increasing temperature as proposed by Brown *et al.* (2013) and use this function along with the ΔVrn1_CL_ defined above, the cool treatment temperature (T) and an exponential coefficient (k = -0.18) to determine the maximum cold vernalization rate parameter is calculated as:

Equation 38.	$\Delta Vrn1Max = {rVrn1}_{CL} / e^{k\times T}$

Finally, this rate is normalized relative to VrnB to give the genotype specific cold vernalization sensitivity coefficient (rVrn1):

Equation 39.	$rVrn1=\Delta Vrn1Max/{rVrnB}_{Veg}$

Figure SC5 shows the derivation of these parameters for three wheat cultivars with contrasting vernalization sensitivity, photoperiod sensitivity, and vernalization by photoperiod interactions (data from Bloomfield *et al.*, 2023). The cultivar ‘Axe’ has very small vernalization and photoperiod sensitivities, with no vernalization and photoperiod interactions. Cultivar Bolac has moderate vernalization and strong photoperiod sensitivities, with no vernalization and photoperiod interactions. Cultivar Wyalkatchem has strong vernalization sensitivity, low photoperiod sensitivity and strong photoperiod and vernalization interactions

#### 1.3 Source code of the CAMP model

The CAMP model was firstly programmed in Python as stand-alone model for deriving model parameters and testing against the data collected from controlled environments and field experiments. It was then implemented in the APSIM NG framework in C# language.

*The source code is freely accessible through the APSIM Use Licence. We provide the following links to the source code now to facilitate the review process.*

The Python code can be found at: [https://github.com/HamishBrownPFR/CAMP](https://checkpoint.url-protection.com/v1/url?o=https%3A//github.com/HamishBrownPFR/CAMP&g=NTk2NWQxNjljZWI2ODFjNg==&h=NDg3YTI5Y2IxMzAwZTVlMGJjMzUyMjY4Y2JhN2ZiOGJhZmJlMzk2YjhmOTU5NWNhZWVkNzUwOGI1NWM0MDFkMw==&p=YzJ1OnBsYW50YW5kZm9vZHJlc2VhcmNoOmM6bzoxOWIxODdkNmM1MDEzMDhkZGFiYWJlZGY5YmE4ZDkyMDp2MTpwOlQ=)

The APSIM NG C# code can be found at: [GitHub - APSIMInitiative/ApsimX](https://checkpoint.url-protection.com/v1/url?o=https%3A//github.com/apsimInitiative/apsimx&g=ZGJhMjI0MjdhZTY3NTM4ZA==&h=NjZlODU2YzAxZWExOTRmZGZmNWY5NGFkYTUxMGU5M2M5OWZmMzEwZmUzZTdmNjYwNTczODdhYjMyZmU0MzhhNA==&p=YzJ1OnBsYW50YW5kZm9vZHJlc2VhcmNoOmM6bzoxOWIxODdkNmM1MDEzMDhkZGFiYWJlZGY5YmE4ZDkyMDp2MTpwOlQ=)

ApsimX is the next generation of APSIM.

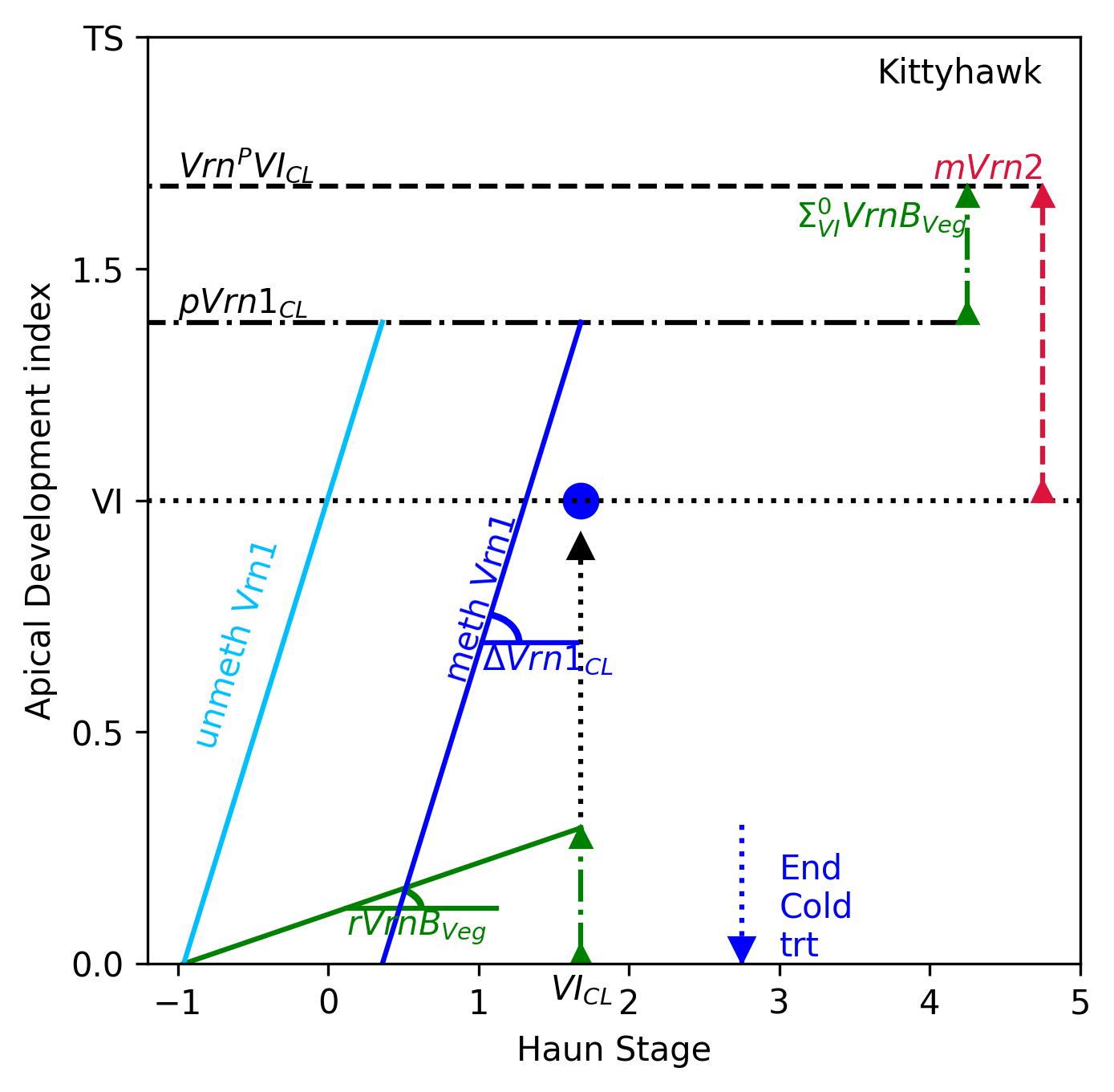

Figure SC4. Plot demonstrating the calculation scheme for deriving *rVrn1* under treatment CL for wheat cultivar Kittyhawk (Bloomfield *et al.*, 2023). *Vrn^P^VI_CL_* is the protagonistic Vrn expression required to achieve VI under long day condition, is equal to the VI threshold (1.0) plus the Vrn needed to offset *mVrn2*. Assuming Vrn3 expression to be negligible before VI under CL (due to presence of Vrn2), the protagonistic gene expression will be from base expression ($\sum_{VI}^{0} {VrnB}_{Veg}$) and the persistent Vrn1 (*pVrn1_CL_*). The same amount of unmethylated Vrn1 must be accumulated before VI. The duration for Vrn1 upregulation by cold is from seed imbibition to VI, and the slope is *△Vrn1_CL_*, which is the rate of cold upregulation of Vrn1 per unit Haun leaf stage. *△Vrn1_CL_* is used to calculate the *rVrn1* in equation 37 &38.

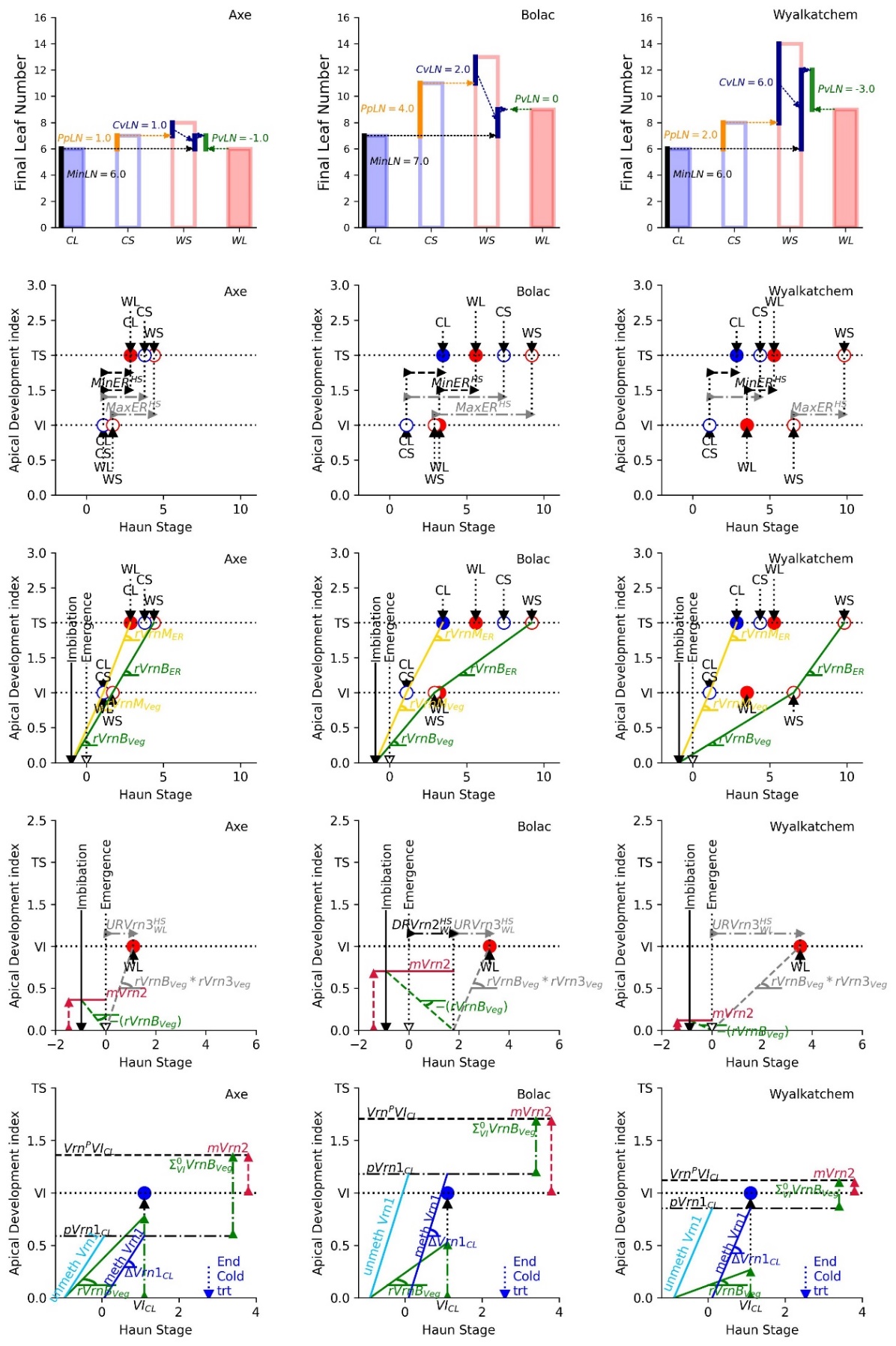

Figure SC5. Mainstem final leaf numbers (top row), minimum and maximum duration of early vegetative stage (2^nd^ row), base and maximum gene expression rates (3^rd^ row), maximum Vrn2 expression (*mVrn2*) (4^th^ row) and cold upregulation of Vrn1 (5^th^ row) of three wheat genotypes (Bloomfield *et al.*, 2023).
